# HCN1 channels in GABAergic amygdalar neurons underpin male-biased aggressive behaviors

**DOI:** 10.1101/2024.12.07.627305

**Authors:** Kaiyuan Li, Zhenggang Zhu, Quan Wang, Lina Pan, Lu Miao, Guochuang Deng, JinRong Wu, Huifang Lou, Ling-Hui Zeng, Yijun Liu, Xiao-ming Li, Shumin Duan, Li Sun, Yan-qin Yu

**Affiliations:** Department of Neurology of Second Affiliated Hospital and School of Brain Science and Brain Medicine, Zhejiang University School of Medicine, Hangzhou 310058, China; Nanhu Brain-computer Interface Institute, Hangzhou 311100, China; Liangzhu Laboratory, MOE Frontier Science Center for Brain Science & Brain-Machine Integration, State Key Laboratory of Brain-machine Intelligence, Zhejiang University, Hangzhou 311121, China; NHC and CAMS Key Laboratory of Medical Neurobiology, Zhejiang University, Hangzhou 310058, China; Key Laboratory of Novel Targets and Drug Study for Neural Repair of Zhejiang Province, School of Medicine, Hangzhou City University, Hangzhou, 310015, China; Institute of Neuroscience, State Key Laboratory of Neuroscience, Chinese Academy of Sciences, Shanghai 200031, China

## Abstract

Aggression behaviors typically vary between sexes, but the molecular mechanisms driving these disparities in neural coding are unclear. We found that aggression selectively activates GABAergic neurons in the posterior substantia innominata (pSI), an extend amygdala region critical for aggressive behaviors in both sexes of mice, with males exhibiting higher neuronal activity during the attack. Utilizing single-nucleus RNA sequencing, we characterized the diverse molecular landscape of pSI neurons, revealing significant differences in ion channels and hormone regulator genes that may underpin sex-specific aggression. Male GABAergic pSI neurons exhibited remarkable hyperexcitability due to increased Ih currents. Strikingly, modulating HCN1 expression not only adjusted this hyperexcitability but also influenced sexual dimorphism in aggression: silencing HCN1 in the GABAergic pSI neurons reduced male aggression, while its overexpression markedly heightened aggression in females. Furthermore, testosterone was shown to intensify aggression by upregulating HCN1 and remodeling pSI circuits. These findings provide detailed sex-specific molecular mechanisms underlying social behaviors.

## Introduction

Sexually dimorphic aggression is a well-documented phenomenon in both animals and humans (*1, 2*). However, both male and female social behaviors can be triggered by similar external or internal factors, suggesting that the underlying mechanisms of aggression may be conserved across the sexes (*3*). Multiple neural circuits governing aggression have been identified, including those that elicit sexually equivalent behaviors— exhibiting similar aggression levels across both sexes upon circuit activation (*4, 5*)—and sexually differentiated, manifesting in distinctly varied aggression levels between males and females following circuit activation (*2, 6–8*). It is plausible that differential activities of these neural circuits across the sexes potentially form the biological basis for observed sex differences in aggression. However, the precise molecular mechanisms gate the neuronal activity levels in these brain regions leading to sex-specific aggressive behaviors remain inadequately explored (*1, 2, 8–10*).

Regarding the molecular basis, hormonal signaling, receptors, and ion channels are closely involved in social behaviors. A notable trend is the recent surge in single cell sequencing studies of key brain nodes in social behavior, which provides molecular insights into how neurons associated with social behaviors express genes differently in males and females. For instance, gonadal steroid hormones through hormone response element to regulate the expression of sex-biased genes, and dynamic effects on neural activity and behavior (*1, 5, 11–18*). Although these molecular differences regulate sexually dimorphic aggressive behaviors, few studies have explored how they impact the encoding of neuronal activity associated with sex-specific aggression.

The posterior substantia innominata (pSI) has been identified to play a central role in modulating aggression in both male and female mice (*4*). Notably, pSI is unidirectionally innervated by VMHvl and regulates sex-biased aggressive behaviors by integrating the excitation-inhibition balance of synaptic inputs from VMHvl (*7*). Interestingly, activation of pSI promotes robust female aggression that is similar to male aggression, while manipulating either the excitatory or inhibitory synaptic inputs of the VMHvl-pSI circuit rarely induce an increase in aggressive behaviors in females (*7*), suggesting that the downstream region pSI may have a mechanism that integrates dynamic excitability to regulate sex-dimorphic aggressive behaviors. One possibility is the presence of a local excitation filter, where the pSI is less excitable, preventing females from exhibiting strong aggressive behaviors. However, the mechanisms underlying the excitability regulation of aggression-controlling regions, including the pSI, remain understudied. In particular, little is known about the roles of specific molecules, such as ion channels and receptors, in mediating the sex-specific intrinsic excitability of aggression-promoting circuits, which may contribute to sex-biased aggressive behaviors.

To further exlore neuronal functions underlying sex-biased aggression, we employed single-nucleus RNA sequencing (snRNA-seq) and multiplex fluorescence in situ hybridization (FISH) to characterize the transcriptional profiles of various cell types in the pSI of both sexes. We found that the excitability of GABAergic pSI neurons, driven by upregulation of the hyperpolarization-activated cyclic nucleotide-gated (HCN) cation channels, encodes male-biased aggression. Strikingly, manipulating HCN1 expression in the GABAergic pSI robustly altered aggressive behaviors in both males and females. Together, our study elucidates the molecular-level sex differences in the pSI, providing valuable insight into the mechanisms underlying the quantitative variations in social behaviors between the males and females.

## Results

### GABAergic pSI neuronal activity encoding male-biased aggressive behavior

To delineate the quantitative differences in aggressive behaviors in mice, we quantified the aggressive behaviors and confirmed that males displayed higher aggression levels with shorter mean attack latency, more attacks, and longer attack durations than females (Extended Data Fig. 1a-h). To unravel the quantitative differences of pSI neurons in sexual dimorphic aggression, we first noticed pSI neurons were selectively activated after aggression in both males and females, indicated by the c-Fos, a genetic marker of neuronal activity (Extended Data Fig. 1i-j). Interestingly, male pSI neurons showed more c-Fos expression compared to females, thus, it is important to examine whether there are different populations (such as transcriptional profiles or electrophysiology-defined neuronal subtypes) of pSI neurons are activated or just the quantitatively differential activation of the same population of pSI neurons. To systematically characterize the transcriptional profiles of single cells in the pSI, we first utilized 10X single-nucleus RNA sequencing (snRNA-seq). After dissecting and digesting the pSI tissue, we profiled 15719 cells (Fig. 1a) and identified 25 distinct cell clusters (Fig. 1b). These clusters can be further classified into 10 GABAergic neuron clusters, 2 glutamatergic neuron clusters, 2 astrocytes clusters, 5 oligodendrocytes clusters, 2 microglia clusters, 1 endothelial cluster (Fig. 1c), and 83.16% of neurons consist of GABAergic neurons, which were classified into different cell type based canonical gene markers (Extended Data Fig. 2b-c). Using multiplex fluorescence in situ hybridization (FISH), we further confirmed that the c-Fos neurons highly co-expressed Slc32a1 (a marker of GABAergic neurons) in both males and females, implying the pSI GABAergic neurons are specifically responsible for aggressive behaviors for both sexes (Extended Data Fig. 3a-b). Interestingly, since the co-expression ratio of Slc32a1 with the c-Fos is even highly than the Camk2α/Thy1 marker that identified previously (*4*), we also characterized the distribution and co-expression of Camk2α/Thy1/Slc32a1 in the pSI (Fig. 1d-e) and found ∼82.52% of Camk2α/Thy1 neurons also expressed the Slc32a1 (Fig. 1f). Using the fiber photometry recording of pSI GABAergic neurons during social aggression in both male and female Vgat-Cre mice (Fig. 1a-b), we found that aggression evoked strong activity in pSI^Vgat^ neurons during aggressive behaviors in both sexes (Fig. 1c), and higher pSI^Vgat^ neuronal dynamics were observed in males than females during the attack (Fig. 1d-g). Although, compared to males, female mice rarely engage in natural aggressive behaviors (Extended Data Fig. 1c-d), robust female aggression was found when pSI^Vgat^ neurons were adequately activated by optogenetics (Extended Data Fig. 3d-f) or pharmacogenetics (Extended Data Fig. 4) methods, suggesting that the insufficient activation of pSI neurons is responsible for the rarity of spontaneous aggressive behaviors in female mice.

**Fig. 1.**
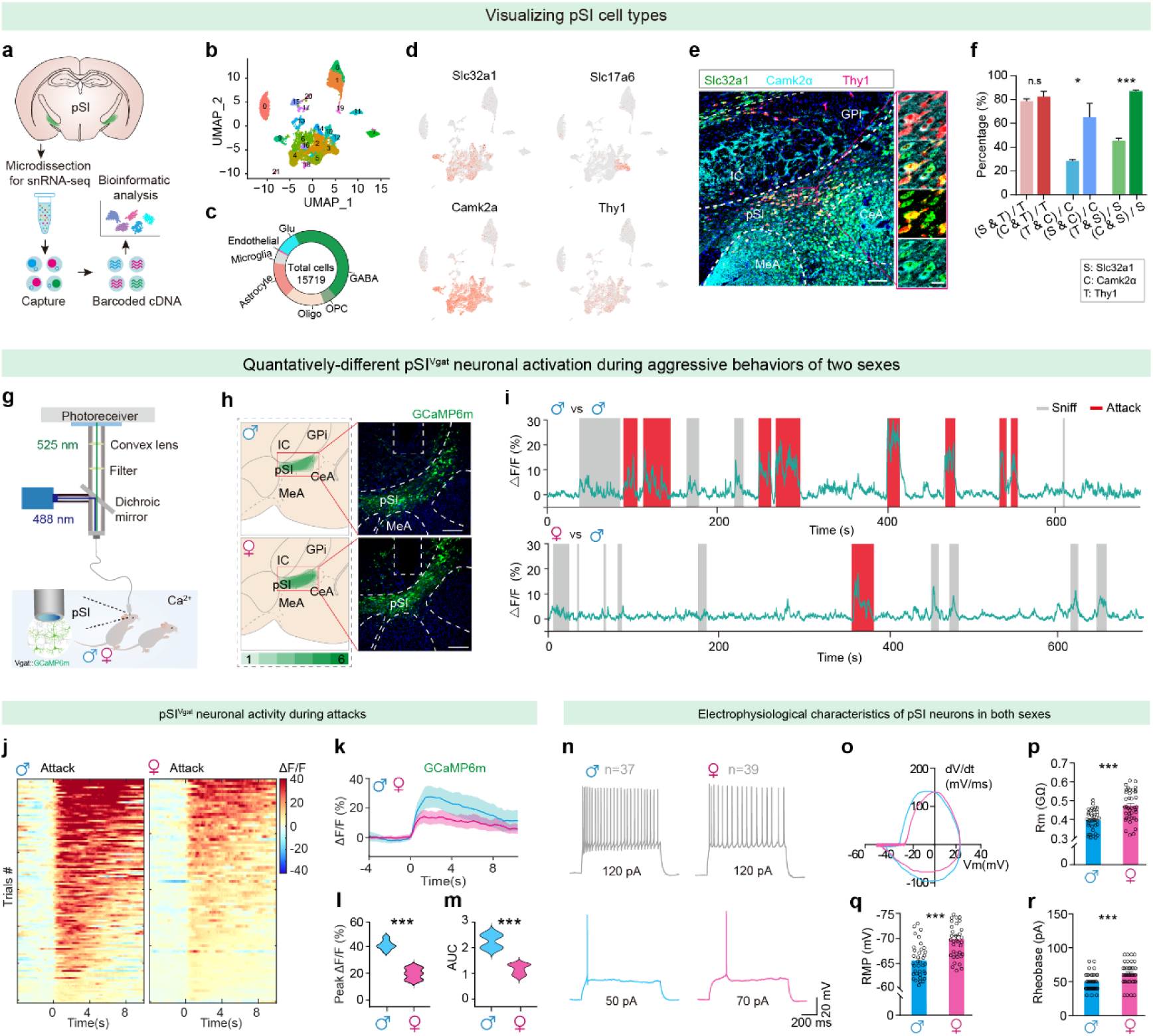
Hyperexcited GABAergic pSI neurons promote aggressive behavior of both sexes. **(a)** Schematic representation of the workflow for pSI microdissection, single-nucleus isolation, cDNA library preparation, sequencing and clustering. Location of pSI microdissections from male and female C57BL/6 mice, mapped onto coronal mouse brain atlas images. **(b)** Two-dimensional UMPA visualization showing the distribution of cells from pSI. **(c)** The proportion of different cell types in the pSI. **(d)** Feature plots showing the expression patterns of cell marker genes in each neuronal cluster. **(e)** Left, a representative image of the expression of Camk2α, Thy1, and Slc32a1 mRNA in pSI in male mice. Scale bars, 200 μm (left). Right, the enlarged images of the expression. Scale bars: 10 μm. (**f**) Co-expression ratio of the Slc32a1 or the Camk2α expression in all Thy1 positive pSI neurons (left). Co-expression ratio of the Thy1 or the Slc32a1 expression in all Camk2α positive pSI neurons (middle). Co-expression ratio of the Thy1 or the Slc32a1 expression in all Slc32a1 positive pSI neurons (right). **(g)** Setup for fiber photometric recording in pSI^Vgat^ neuronal activity during social behaviors. **(h)** Representative pSI^Vgat^ neurons GCamp6m signals of attack behavior recorded during encounters in males (upper) and females (lower) exposed to a male. Scale bars, 200 μm. **(i)** Representative pSI^Vgat^ GCamp6m signals when males (upper) and females (lower) are exposed to a male. **(j)** Heatmap of GCamp6m ΔF/F signals (male: upper, female: lower) of pSI^Vgat^ neurons in attacks, n = 121 trials in males, n = 92 trials in females. **(k)** Summary of pSI^Vgat^ neuronal activity (ΔF/F) recording from male and female mice during the attack. **(l-m)** peak ΔF/F (f), and area under the curve (AUC, g) of the pSI^Vgat^ neuronal activity in males and females. **(n)** Two distinct representative spiking patterns were observed in pSI neurons from both males (blue, n = 37) and females (magenta, n = 39) after a pulse train was intracellularly injected. The injection current each train lasted for a duration of 1 second. **(o)** The first derivative of the membrane voltage (dV/dt) vs membrane voltage (Vm), for pSI-PAG neurons in male (blue line) and female (magenta line) mice. Representative phase-plot showing the rate of change of the action potential during all 3 phases of the spike: depolarization, repolarization, and refractory slope. **(p-r)** The membrane resistance (Rm, p), resting membrane potential (RMP, q), and rheobase (r) of all the recorded male and female pSI neurons. In (f) and (p-r): Mann-Whitney U test; (l-m): Unpaired t-test with Welch’s correction; *P < 0.05, ***P<0.001, n.s, not significant. For all panels, data are shown as the mean ± SEM. Statistical details are available in the supplementary table S1.

To test whether sexual differences of aggression could be correlated to intrinsic electrophysiological properties of pSI neurons, we systematically compared the intrinsic properties of pSI neurons, including membrane resistances (Rm), resting membrane potential (RMP), and rheobase, by whole-cell patch clamp recordings. We found a significant difference in electrophysiological properties between males (37 neurons from 5 mice) and females (39 neurons from 5 mice), such as the Rm (Fig. 1j) being larger in females than males, while the RMP (Fig. 1k) was more depolarized in males and the rheobase (Fig. 1l) was lower in males than in females. Thus, male pSI neurons are fired with a smaller depolarization indicating pSI is more excitable in males. Together, these electrophysiology-defined neuronal properties of pSI neurons suggest that rarely naturally robust aggression in female mice may be linked to specific molecular substrates that gate the threshold of pSI^Vgat^ neuronal activation for trigger aggression.

### Sex-biased gene expression and intrinsic excitability of GABAergic pSI neurons

The difficulty in activating pSI neurons for eliciting natural female aggression implies that the intrinsic molecular properties of the pSI neurons might differ between males and females. As a relatively unexplored region, the sex-biased gene expression of pSI neurons is unknown. To better understand the intrinsic molecular properties of pSI neurons in two sexes, we profiled the gene expression of pSI neurons with snRNA-seq. Although no difference was found in the types of clusters of pSI cells between the two sexes (Fig. 2a and Extended Data Fig. 2a), a comparison of gene expression levels in pSI neurons revealed that several hundred genes are expressed differently in males than in females (Fig. 2c). To characterize neuronal cell types in detail, we re-clustered the neuronal population identified in the initial analysis. From this, we identified 13 discrete neuronal clusters (Fig. 2a). This revealed that the GABAergic neuron marker gene Slc32a1 was expressed in the majority of neuronal clusters, and Slc17a6-positive glutamatergic neurons were mainly expressed in clusters 2, 12 and 13, and significant differences in gene expression were observed between GABAergic and glutamatergic neurons (Extended Data Fig. 2d). For instance, genes that regulate neuronal excitability, such as ion channels, the glutamate receptor, calmodulin-dependent protein kinase, are expressed more in the GABAergic neurons than the glutamatergic neurons of the pSI (Extended Data Fig. 2e). Furthermore, the proportions of GABAergic and glutamatergic neurons in the pSI were confirmed with the fluorescence *in situ* hybridization (FISH) (Fig. 2b), consistent with the FISH results from the Allen Atlas. To characterize molecular differences in GABAergic neurons to the sexually dimorphic aggressive behavior, we identified several sex-differentiated genes in the GABAergic pSI neurons (Fig. 2d and Extended Data Fig. 2f-g). Subsequently, we performed differential gene GO enrichment pathway analysis on snRNA-seq data from male and female mice. The results revealed significant sex differences in multiple biological processes. Specifically, male mice showed enriched gene expression in pathways such as calcium signaling, dopamine signaling, 5-HT signaling, and glutamatergic synapses, whereas female mice exhibited less enrichment in these pathways (Fig. 2e). These findings suggest that genes related to hormone or its receptor and ion channels are abundantly located in pSI GABAergic neurons and might be involved in the sex-specific aggressive behaviors in male and female mice.

**Fig. 2.**
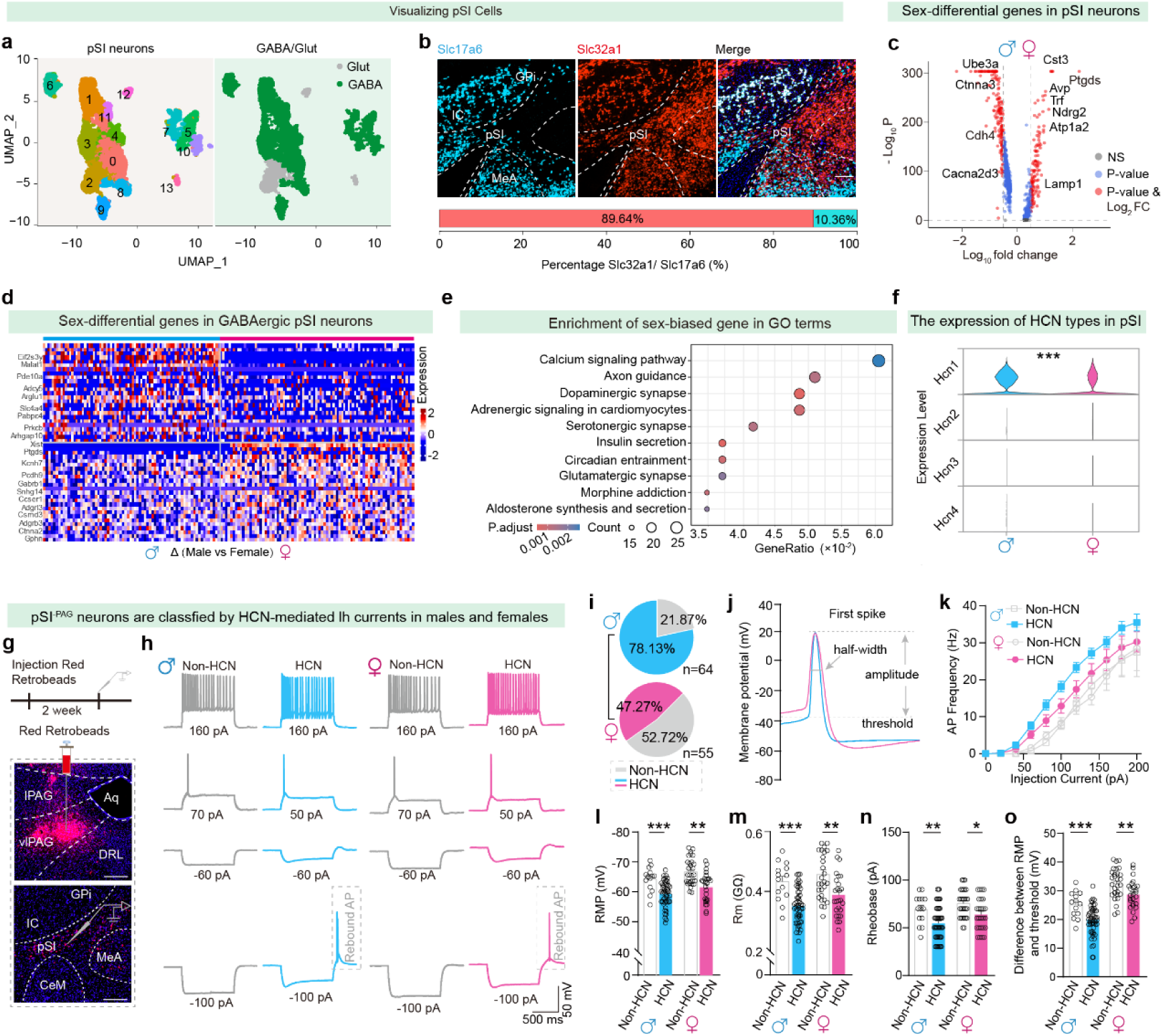
Molecular profiles and electrophysiological properties of GABAergic pSI neurons between males and females. **(a)** Two-dimensional UMPA visualization showing the cluster distribution of all neurons from males and females (left), GABAergic and glutamatergic cells, and merged cells of two sexes (right) among major pSI neuron types. **(b)** Representative pSI FISH images showing the overlay of Slc17a6^+^ cells (cyan) and Slc32a1^+^ cells (red). Scale bar, 200 μm. Lower, the percentage of Slc32a1^+^ cells (red) and Slc17a6^+^ cells (cyan) from all pSI neurons. **(d)** The volcano plot of differential genes of males versus females in the pSI. **(d)** Differentially expressed genes in GABAergic pSI neurons between males and female. **(e)** Enrichment of sex-biased gene in GO terms between males and female. **(f)** Distribution of the cyclic nucleotide regulated channel family in the male and female pSI from the snRNA-seq. **(g)** Representative images of retrobeads tracking and experimental scheme, showing infections of retrobeads in the PAG (Upper), and retrograde labeling of pSI^-PAG^ neurons (Lower). Scale bars represent 200 µm (upper). **(h)** Two distinct representative spiking patterns (with HCN or without HCN mediated rebound depolarization) were observed in pSI^-PAG^ neurons from both males (gray, n = 14 non-HCN neurons; blue, n = 50 HCN neurons) and females (gray, n = 16 non-HCN neurons; magenta, n = 29 HCN neurons) after a pulse train was intracellularly injected. The injection current ranged from - 100 pA to 160 pA in 10 pA steps, and each train lasted for a duration of 1 second. **(i)** Relative proportions of HCN neurons and non-HCN neurons were recorded from pSI^-PAG^ neurons in male and female mice (n = 64/55 neurons from 10 mice in the male and female group). **(j)** The first spike elicited in an example neuron was processed to calculate the electrophysiological parameters of the action potential: threshold, amplitude, and half-width duration. **(k)** The spike frequency of male and female pSI^-PAG^ neurons during the step current injections, which triggered more spikes in male than female pSI^-PAG^ neurons. **(l-o)** The resting membrane potential (RMP, l), membrane resistance (Rm, m), Rheobase (n), and the difference between RMP and AP threshold (o) were compared between male and female pSI^-PAG^ neurons. In (c) to (f): Wilcoxon rank-sum test; (k): two-way ANOVA followed by post hoc Šidák’s multiple comparisons tests; (l) to (o): Mann-Whitney U test. *P < 0.05, **P < 0.01, ***P < 0.001. For all panels, data are shown as the mean ± SEM. The electrophysiological properties of pSI^-PAG^ neurons are detailed statistically in supplementary table S1. Statistical details of the snRNA-seq are available in supplementary table S2.

Single-cell transcriptomic analysis of the pSI neurons suggests the sex-biased gene profiles related to neuronal excitability may vary between the two sexes. One hypothesis is that these pSI neurons with such profiles might be less intrinsically excitable in females under natural physiological conditions so that female aggression occurs naturally at a much lower threshold than in males. To investigate whether the intrinsic electrophysiological properties of GABAergic pSI neurons differ in the two sexes, we injected retrobeads into the periaqueductal gray (PAG, a downstream region densely projected by GABAergic pSI neurons that dominates aggressive behavior in mice, Extended Data Fig. 5) and performed the whole-cell patch-clamp recordings from the aggression-related pSI^-PAG^ neurons (Fig. 2g-o) in acute brain slices from males (64 neurons from 5 mice) and females (55 neurons from 6 mice) (Fig. 2i). Strikingly, the co-expression ratio of retrobeads-expressing pSI neurons with the Slc32a1 is highly in Vgat-venus mice (Extended Data Fig. 5h). Ion channels are known to modulate neuronal excitability which can lead to neuronal hyperexcitability (*19–26*). We found that HCN1 channels (a subtype of hyperpolarization-activated cyclic nucleotide-gated nonselective cation channel) were highly specifically distributed in the pSI and differentially expressed in pSI neurons between the two sexes (Fig. 2f), which constitute the molecular substrate of a persistent potassium and sodium current (also known as Ih), and the second messenger molecule cAMP can modulate the kinetic properties of HCN channels.

To systematically investigate the differences in intrinsic properties of pSI^-PAG^ neurons, we compared membrane properties and firing patterns related to the action potentials (AP) while recording pSI^-PAG^ neurons between males and females (Fig. 2j-o and Extended Data Fig. 6). Notably, we found that male pSI^-PAG^ neurons had ∼78.13% shown a depolarization voltage “sag”, which was higher than that in females (only about 47.27%) (Fig. 2h-i). These sag potentials in current-clamp recordings indicate the activation of Ih currents and the molecular substrate of the HCN channel expressed in these neurons (*27, 28*). Specifically, pSI^-PAG^ neurons showed a bimodal distribution of all firing patterns between HCN and non-HCN neurons in both males and females. For example, HCN neurons also exhibited lower step-current evoked high-frequency firing activity than those of non-HCN neurons (Fig. 2k). The resting membrane potential (RMP) of HCN neurons was more depolarized than non-HCN neurons (Fig.2l) while the membrane resistances (Rm, Fig. 2m) being larger in non-HCN neurons than HCN neurons in both males and females. The minimum current injection needed to produce an AP, known as the rheobase, was lower in HCN neurons than in non-HCN neurons (Fig. 2n), indicating that APs are fired more frequently after smaller depolarizations in HCN neurons. The difference between RMP and threshold that relates to the action potentials (APs) was lower in HCN neurons compared to non-HCN neurons (Fig. 2o). More importantly, when we compared the electrophysiological properties of HCN neurons between males and females, HCN neurons in females exhibited higher Rm and higher firing threshold, while HCN neurons in males displayed shorter-latency APs and higher APs firing frequency (Fig. 2l-o, and Table 1). Moreover, we compared 17 electrophysiological features in recorded pSI^-PAG^ neurons from both males and females and categorized by whether they expressed the HCN or not (Table. 1). The ability of the HCN channel to predict the bimodal distribution of electrophysical properties of the pSI^-PAG^ neurons suggests the crucial role of the HCN channel in neuronal hyperexcitability in the pSI of mice.

### Sexually disparate HCN-mediated intrinsic excitability of pSI^-PAG^ neurons

To better characterize the HCN-mediated electrophysical hyperexcitability of pSI neurons of both sexes, we directly measured the amplitude of Ih currents in pSI^-PAG^ neurons by 1-s hyperpolarizing voltage steps (− 20 mV) held at − 40 mV using whole-cell patch clamp recording in males and females. We found that the Ih current is higher in pSI^-PAG^ neurons of males than in females and can be effectively blocked by HCN channel blockers (Fig. 3a-b). However, no significant differences in Na^+^ and K^+^ currents were observed in pSI^-PAG^ neurons of female and male mice (Extended Data Fig. 6c-e). To further confirm sexually differential gene expression in GABAergic pSI neurons, we performed fluorescent in situ hybridization (FISH) to examine the co-expression of the Slc32a1 and HCN1 genes in pSI (Fig. 3c). We found GABAergic pSI neurons highly co-expressing the molecular markers of HCN1, and HCN1 expressed more abundantly in the pSI of males than females (Fig. 3d-f). Application of an extracellular HCN antagonist, ZD7288, significantly altered the intrinsic electrophysiological properties of HCN neurons, such as hyperpolarized the RMP, and increased the Rm of HCN neurons, but did not significantly alter non-HCN neurons in males (Fig. 3g-j and Extended Data Fig. 6a-b). These suggest the HCN channel expression determines the neuronal hyperexcitability of the pSI^-PAG^ neurons. Additionally, our further analysis of HCN1-related genes in the GO term from the snRNA-seq data revealed that many genes associated with neuronal membrane potential, synaptic transmission, and the regulation of synapse organization could be enriched in HCN1-positive neurons (Fig. 3k), implying the potential molecular repertories for HCN1 expression that may determine the neuronal hyperexcitability by altering the intracellular environment. Altogether, multiple key electrophysiological properties in the pSI^-PAG^ neurons implied a higher intrinsic neuronal excitability of the pSI neurons in males than in females, which is mediated by the expression of the HCN1.

**Fig. 3.**
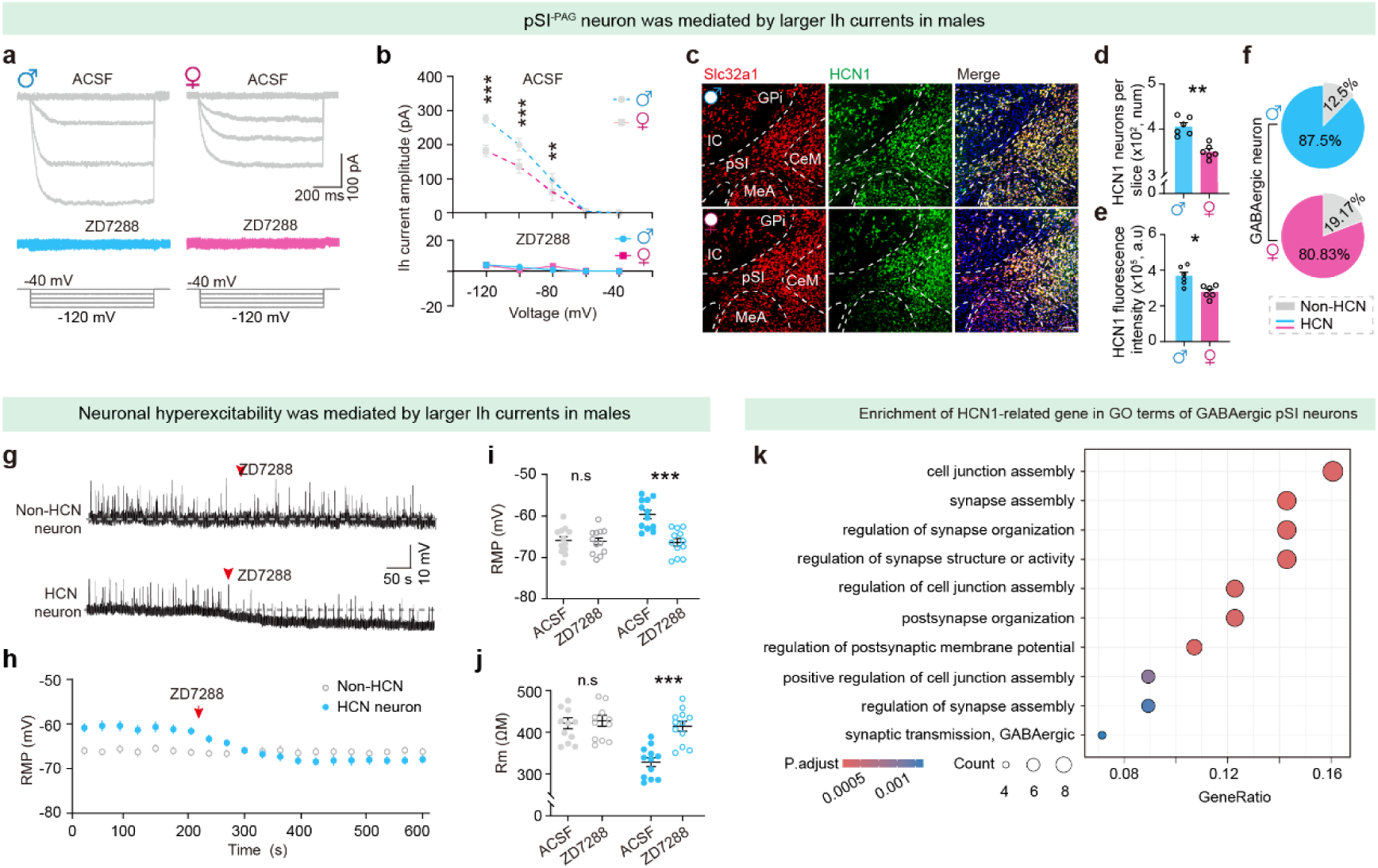
Increased HCN currents (Ih) leads to hyperexcitability of pSI^-PAG^ pSI neurons in male mice. **(a-b)** Comparison of Ih current between male and female pSI^-PAG^ neurons during the step voltage injections before and after ZD7288. Whole-cell patch clamp recording neuron resting membrane potential holding at − 40 mV in response to 1 s voltage steps (0 to − 80 mV, − 20 mV step). All neurons recorded in ACSF including TTX 1 μM, 4-AP 100 μM, CNQX 40 μM, APV 100 μM, bicuculine 100 μM, and BaCl_2_ 100 μM. **(c)** Representative pSI FISH images showing the overlay of Slc32a1^+^ cells (red) with HCN1^+^ cells (green) in the pSI of C57BL/6J male (left) and female (right) mice. Scale bar, 100 µm. **(d-f)** Cell number of HCN1^+^ neurons (d), the fluorescence intensity of HCN1^+^ neurons (e), and the percentage of Slc32a1^+^ cells (red) that overlap with HCN1^+^ cells (f) in the pSI of male and female mice (n = 6 mice in each group). **(g)** Representative voltage trace of HCN and non-HCN neurons across time. **(h)** Averaged resting membrane potential across time of the pSI^-PAG^ neurons before or after the perfusion of ZD7288 (100 μM). **(i-j)** Comparison of resting membrane potential (RMP, i), and the membrane resistance (Rm, j) between male and female pSI^-PAG^ neurons during the step current injections. (**k**) GO enrichment analysis of the HCN1 positive cluster in the GABAergic pSI neurons. In (b): Two-way ANOVA followed by *Post-hoc* Šidák’s multiple comparisons; (d) to (e): Mann-Whitney U test; (i) and (j): paired t test; (k): Wilcoxon rank-sum test; *P < 0.05, **P < 0.01, ***P < 0.001, n.s, not significant. For all panels, data are shown as the mean ± SEM. The electrophysiological properties of pSI^-PAG^ neurons are detailed statistically in supplementary table S1. Statistical details of the snRNA-seq are available in supplementary table S2.

### HCN channel-regulated GABAergic pSI neuronal hyperactivity drives aggression in males

Neuronal hyperactivity mediated by the HCN1 channel in the pSI potentially facilitates aggression in mice. To further test the functional role of HCN1 channels in the pSI for aggression, we pharmacologically block the HCN channels in the pSI by bilaterally implanting a cannula above the pSI and infusing the HCN channel blockers into the male mice. The HCN blockers ZD7288, zatebradine, and ivabradine, but not the vehicle control, hyperpolarized the resting membrane potential and decreased the AP firing frequency in pSI neurons (Fig. 4a-c and Extended Data Fig. 7a-f). They also increased the rebound AP latency and significantly reduced the sag amplitude (Fig. 4d-f). Two weeks after cannula implantation surgery, the HCN blockers were infused into the pSI of the singly housed male mice (Fig. 4g-h). Interestingly, local blocking of HCN channels by the injection of these HCN blockers did not significantly affect mice locomotion during the open field test (Extended Data Fig. 7g-i). However, the HCN blockers— ZD7288, zatebradine, and ivabradine, but not the control vehicle, significantly reduced male aggressive behaviors during the aggression test (Fig. 4i and Extended Data Fig. 7j-k). Notably, chemicals that cross the blood-brain barrier (BBB), such as ZD7288, but not ivabradine or zatebradine, could have greater therapeutic potential. We further intraperitoneally administered the HCN blockers-ZD7288, zatebradine, and ivabradine in a dosed manner (0.2, 1, 2, and 5 mg/kg) and tested the aggressive behaviors in singly housed male mice. We observed that ZD7288 significantly reduced male aggression during the aggression test (Fig. 4j). However, intraperitoneally administered either ivabradine or zatebradine or the control group did not significantly affect male aggression (Extended Data Fig. 7l-m). This suggests the pSI^Vgat^ neuronal hyperexcitability was mediated by the HCN channel. cAMP signaling has been shown to activate the HCN channels and mediate Ih currents amplification, thus, pharmacological activation of the HCN channels through cAMP signaling in the pSI may affect the aggressive behaviors (*29–31*). We found that injecting a cAMP activator-forskolin, and the vehicle control into the pSI of male mice that exhibited minimal natural inter-male aggression (Fig. 4k) significantly enhanced the events and duration of male aggressive behaviors, as compared to the control vehicle (Fig. 4l). Together, these pharmacological activations or suppression of the HCN channels of the pSI in aggression supported the hypothesis that HCN1 channels could affect aggression by regulating the neuronal activation in the pSI (Fig. 4m).

**Fig. 4.**
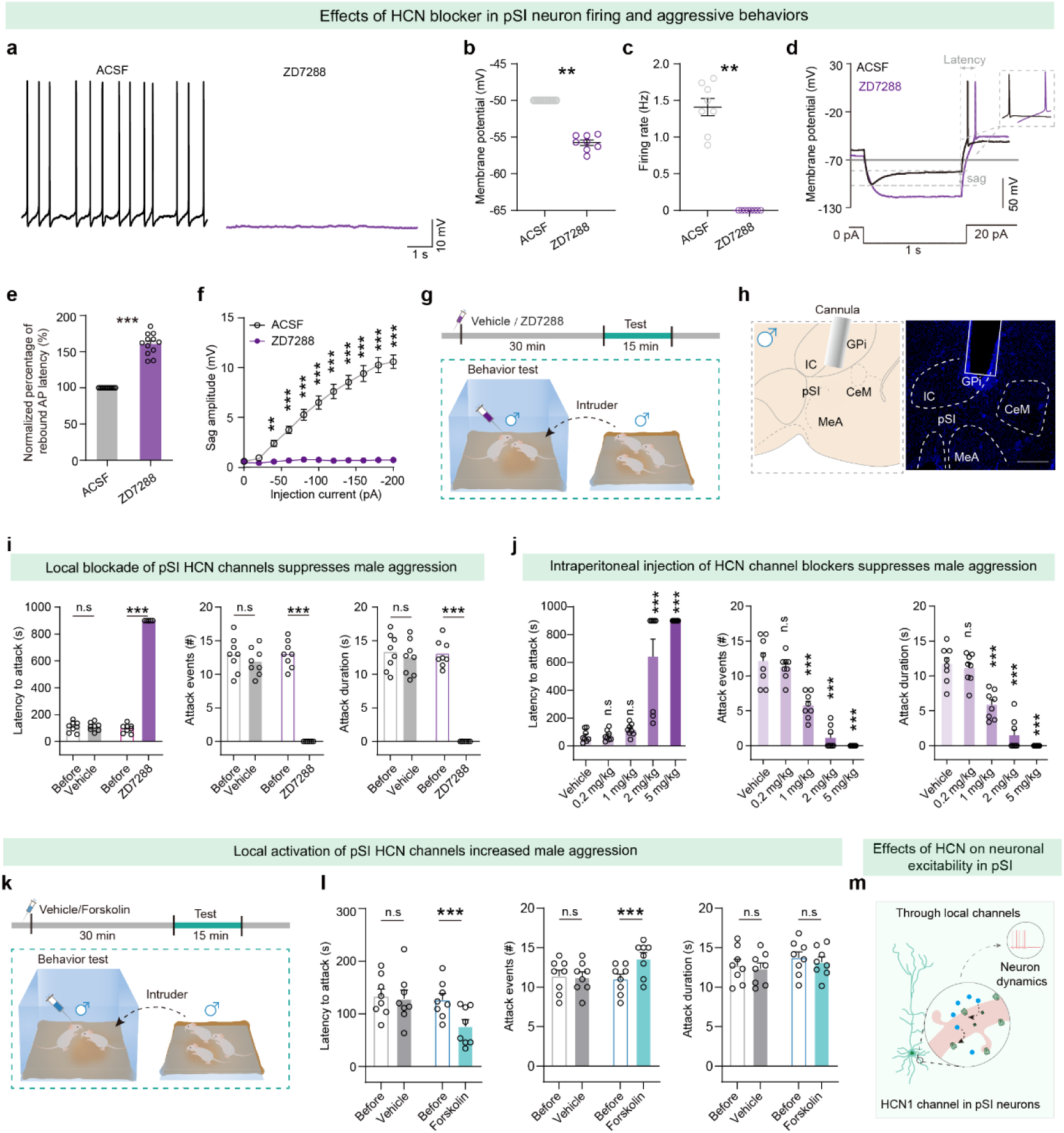
Pharmacological manipulation of pSI HCN1 channels alters aggressive behaviors. **(a-c)** Example electrophysiological traces, RMP (b), and firing rate (c) of pSI neurons with pharmacological inhibition of HCN1 channels in the pSI by ZD7288 (n=12 neurons). (**d**) Sample traces of the spike in rebound burst and hyperpolarization-activated depolarization “sag” were recorded from pSI neurons in male mice treated with ACSF or HCN1 blocker-ZD7288 (100 μM). (**e-f**) Summarized data of the normalized percentage rebound AP latency (e) and hyperpolarization-activated depolarization “sag” amplitude (f) from pSI neurons in male mice treated with control or HCN1 blocker (ZD7288, n=12 neurons). (**g**) The behavioral paradigm of attack test for manipulation of pSI HCN1 channels with the HCN ion channel gated blockers-ZD7288 infused into the pSI in male mice. **(h)** Histology, example trajectory trace showing the inhibition of HCN1 channels with pharmacological inhibition using vehicle control or ZD7288 in the pSI (left, n = 8 mice per group). Scale bar, 500 µm. (**i**) Effects of pharmacological inhibition of HCN1 channels (ZD7288 200 μM, 300 nl per side) in the pSI for aggression. The attack latency, attack events, and attack duration of male mice toward a socially housed male intruder in its home cage (n = 8 mice in each group) were analyzed. For detailed statistical information, see Table S2. (**j**) Effects of pharmacological inhibition of HCN1 channels (ZD7288 at the diverse doses of 0.2 mg/kg, 1 mg/kg, 2 mg/kg, 5 mg/kg in the manner of intraperitoneal injection) for aggression. The attack latency, attack events, and attack duration of male mice toward a socially housed male intruder in its home cage (n = 8 mice/group) were analyzed. (**k**) The behavioral paradigm of attack test for manipulation of pSI HCN1 channels with the HCN ion channel agonist-forskolin infused into the pSI in male mice. (**l**) Effects of pharmacological activation of the pSI HCN1 channels by applying adenosine cyclase agonist (Forskolin, 200 μM; 300 nl) in the pSI for aggression. The first attack latency, attack events, and attack duration of male mice (n = 8 mice in each group) were analyzed. **(m)** Possible mechanisms by which HCN1 channels regulate neuronal activity in pSI. In (b), (c), (e): Wilcoxon matched-pairs signed rank test; (f): two-way ANOVA followed by post hoc Šidák’s multiple comparisons tests; (i) and (l): paired t test and unpaired t test with Welch’s correction; (j): One-way ANOVA followed by Holm-Šídák’s multiple comparisons test; Statistical details are available in the supplementary table S1.

### Disruption of HCN1 genes in GABAergic pSI diminishes hyperactive male aggression

Observed fewer HCN1 expressions of GABAergic pSI neurons in females versus males implies that the regulation of HCN1 expression could be vital to the aggression of mice. To validate this, we used CRISPR/Cas9-mediated disruption of HCN1 genes of pSI^Vgat^ neurons in male mice, with an AAV vectors serotype 2/9 (rAAV_2/9_) based CRISPR/Cas9 system expressing sgRNA with AAV2/9-U6-spsgRNA (HCN1)-U6 spsgRNA (HCN1)-CMV-EGFP-bGH to target HCN1 mRNA, and it was infused bilaterally into the pSI of Vgat-Cre::Cas9 mice (Fig. 5a). The control mice received the same viral vector carrying the fluorophore but not sgHCN1 (rAAV2/9-U6-spsgRNA(scramble)-U6-spsgRNA(scramble)-CMV-EGFP-bGH) (Fig. 5a). After 8 weeks of viral expression, we found that the expression of HCN1 genes in the CRISPR/Cas9-mediated disruption mice was dramatically reduced compared with mice injected with rAAV2/9-U6-spsgRNA(scramble)-U6-spsgRNA(scramble)-CMV-EGFP-bGH (Extended Data Fig. 8d-f). Whole-cell recording from individual EGFP-labeled GABAergic pSI neurons showed reduced Ih currents amplitude, hyperpolarized RMP, and increased rheobase after the knockdown of HCN1 genes (Fig. 5b-e). These results demonstrated that the knockdown of pSI HCN1 genes is effective in downregulation of the HCN1 gene expression and largely impaired the neuronal excitability of the pSI neurons in male mice.

**Fig. 5.**
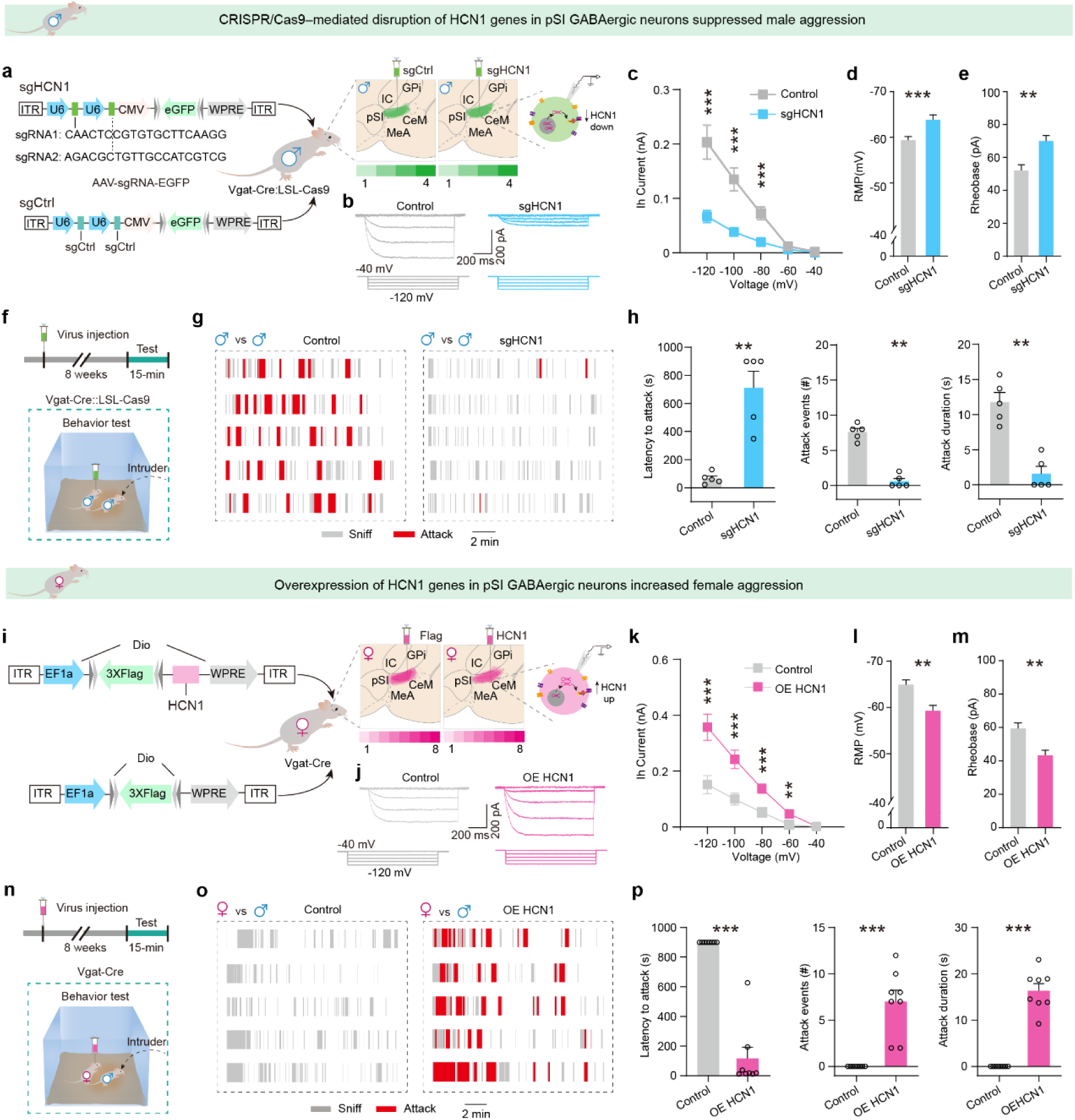
CRISPR/Cas9–mediated disruption and overexpression of HCN1 genes in GABAergic pSI neurons alters the sexual dimorphic aggressive behaviors. (**a**) Design of CRISPR/Cas9– mediated disruption of HCN1 genes by using the AAV sgControl, AAV-hsyn-Cre, AAV sgHCN1 vector design, and bilateral viral infection of pSI neurons in Rosa-LsL-Cas9 male mice. Left, the schematic of the viral injection strategy for the AAV_2/9_-U6-spsgRNA (HCN1)-U6-spsgRNA (HCN1)-CMV-EGFP-bGH (Green) or rAAV_2/9_-U6-spsgRNA(scramble)-U6-spsgRNA(scramble)-CMV-EGFP-bGH (Green) mixture of AAV-hsyn-Cre-PA in the pSI of Rosa-LsL-Cas9 male mice. Upper right, Experimental design for examining the AP firing patterns in the sgControl and sgHCN1 neurons. (**b**) Representative traces of Ih currents were recorded from sgControl (gray) and sgHCN1 (blue) groups. Whole-cell patch clamp recording neuron resting membrane potential holding at − 40 mV in response to 1 s voltage steps (0 to − 80 mV, − 20 mV step). All neurons recorded in ACSF including TTX 1 μM, 4-AP 100 μM, CNQX 40 μM, APV 100 μM, bicuculine 100 μM, and BaCl_2_ 100 μM. (**c-e**) Comparison of the current amplitude (pA, c) of Ih, the RMP (mV, d), and the rheobase (pA, e) between the sgControl and sgHCN1 group. Statistical data of the current amplitude (pA) of Ih is plotted against the voltage step in pSI neurons from the sgControl and sgHCN1 groups. (n = 12 neurons from three mice in each group). (**f**) Behavioral paradigm shows a singly housed Rosa-LsL-Cas9 male mouse in its home cage encountering a socially housed male intruder. (**g**) Raster plots of attack behavior recorded during 15-minute encounters in the sgControl and sgHCN1 groups (red blocks indicate the duration of episodes of attack, and gray blocks indicate the duration of episodes of sniff). (**h**) Latency to the first attack, attack events, and attack duration at 8 weeks after the viral injection, as illustrated in (a). (**i**) Schematic of AAV HCN1, AAV Flag vector design, and bilateral viral infection of pSI neurons in Vgat-Cre female mice. Experimental design and viral injection strategy for the expression of AAV-Flag-HCN1 or AAV-Flag in pSI^Vgat^ neurons in Vgat-Cre female mice and the subsequent behavioral test. Left, schematic of the viral injection strategy for the rAAV_2/9_-EF1α-Dio-3XFlag-HCN1-WPRES or rAAV_2/9_-EF1α-Dio-3XFlag-WPRES in the pSI of Vgat-Cre female mice. Right, representative slices expressing anti-Flag positive neurons in pSI^Vgat^ neurons. Patch clamp recorded cells were labeled by the virus, showing AP firing patterns in the control and overexpression of HCN1 neurons. (**j**) Representative traces of Ih currents were recorded from control (gray) and overexpression of HCN1(magenta) groups. Whole-cell patch clamp recording neuron resting membrane potential holding at − 40 mV in response to 1 s voltage steps (0 to − 80 mV, − 20 mV step). All neurons recorded in ACSF including TTX 1 μM, 4-AP 100 μM, CNQX 40 μM, APV 100 μM, bicuculine 100 μM, and BaCl_2_ 100 μM. (**k-m**) Comparison of the current amplitude (pA, k) of Ih, the RMP (mV, l), and the rheobase (pA, m) between the control and overexpression of HCN1 group. Statistical data of the current amplitude (pA) of Ih is plotted against the voltage step in pSI from control and overexpression of HCN1 groups. (n = 12 neurons from three mice in each group). **(n)** The behavioral paradigm shows a singly housed male Vgat-Cre female mouse in its home cage encountering a socially housed male intruder. (**o**) Raster plots of attack behavior recorded during 15-minute encounters in the control and overexpression of HCN1 groups (red blocks indicate the duration of episodes of attack, gray blocks indicate the duration of episodes of sniff). (**p**) Latency to the first attack, attack events, and attack duration at 8 weeks after injection of the virus. Data are mean ± SEM; (c), (d), (k) and (l): Mann-Whitney U test; (e) and (m): two-way ANOVA followed by post-hoc Šidák’s multiple comparisons; (h) and (p): Mann-Whitney U test; *P < 0.05, **P < 0.01, ***P < 0.001. For all panels, data are shown as the mean ± SEM. Statistical details are available in the supplementary table S1.

Additionally, eight weeks after the virus injection, we performed the open field test and a resident-intruder assay to assess the necessary role of HCN1 genes in pSI neurons in locomotion (Extended Data Fig. 8g-h) and aggression (Extended Data Fig. 8i-j). Notably, CRISPR/Cas9-mediated deactivation of HCN1 genes in pSI neurons (Extended Data Fig. 8I-J), as well as in pSI^Vgat^ neurons (Fig. 5f-h) was sufficient to suppress male aggression by largely increasing the first attack latency and reducing attack events and duration (Fig. 5f-h) but did not affect the locomotion during the open field test in these animals (Extended Data Fig. 8g-h).

### Overexpressing HCN1 genes in GABAergic pSI remarkably enhanced female aggression

To further investigate the role of HCN1 channels in triggering aggression in both sexes, we next examined whether overexpression of HCN1 in pSI^Vgat^ neurons could affect aggressive behaviors in females (Fig. 5i). We designed AAV vectors to overexpress HCN1 mRNA, and the packaged virus was infused bilaterally into the pSI of Vgat-Cre mice (rAAV-EF1α-Dio-3Xflag-HCN1 injected mice). In control rAAV-EF1α-Dio-3Xflag injected mice, the injection site received the same viral vector carrying the 3Xflag but not HCN1 (Fig. 5i). Compared with mice injected with the control virus of expressing rAAV-EF1α-Dio-3XFlag, expression of HCN1 was largely increased in the pSI of rAAV-EF1α-Dio-3XFlag-HCN1 mice (Extended Data Fig. 9a-c). We then performed whole-cell recording from pSI^Vgat^ neurons at 8 weeks after virus injection (rAAV-EF1α-Dio-3XFlag-HCN1 and rAAV-EF1α-Dio-EGFP) and found that the GABAergic pSI neurons with overexpression of HCN1 gene showed significantly increased Ih currents amplitude, depolarized RMP, and decreased rheobase when compared with those of the control group (Fig. 5j-m).

Eight weeks after the virus injection, we performed an aggression test in these rAAV-EF1α-Dio-3XFlag-HCN1 mice and control mice. Remarkably, overexpression of HCN1 genes in pSI^Vgat^ neurons did not significantly affect mice locomotion during the open field test but was sufficient to enhance the attack events and duration in female mice (Fig. 5n-p and Extended Data Fig. 9d-e). Together, we confirmed that selective upregulation of the expression of HCN1 genes could diminish the sexual dimorphism of aggression by largely increasing female aggressive behaviors.

### Testosterone regulates male-biased aggression through the HCN channel in GABAergic pSI neurons

Sex hormones, like testosterone, were found to differ between males and females (Fig. 6a) and are known to play a critical role in shaping aggression during development stages (*12, 13, 32*). For instance, testosterone can masculinize the aggression circuitry and lead to a high level of aggression in males during the development stages. However, the role of these sex hormones in maintaining a high level of aggression in adult animals is less well-known. The disparate neuronal excitability of the pSI circuit potentially suggests a critical entry point for understanding the hormonal control of aggression through pSI in mice (see Figs 1-5).

**Fig. 6.**
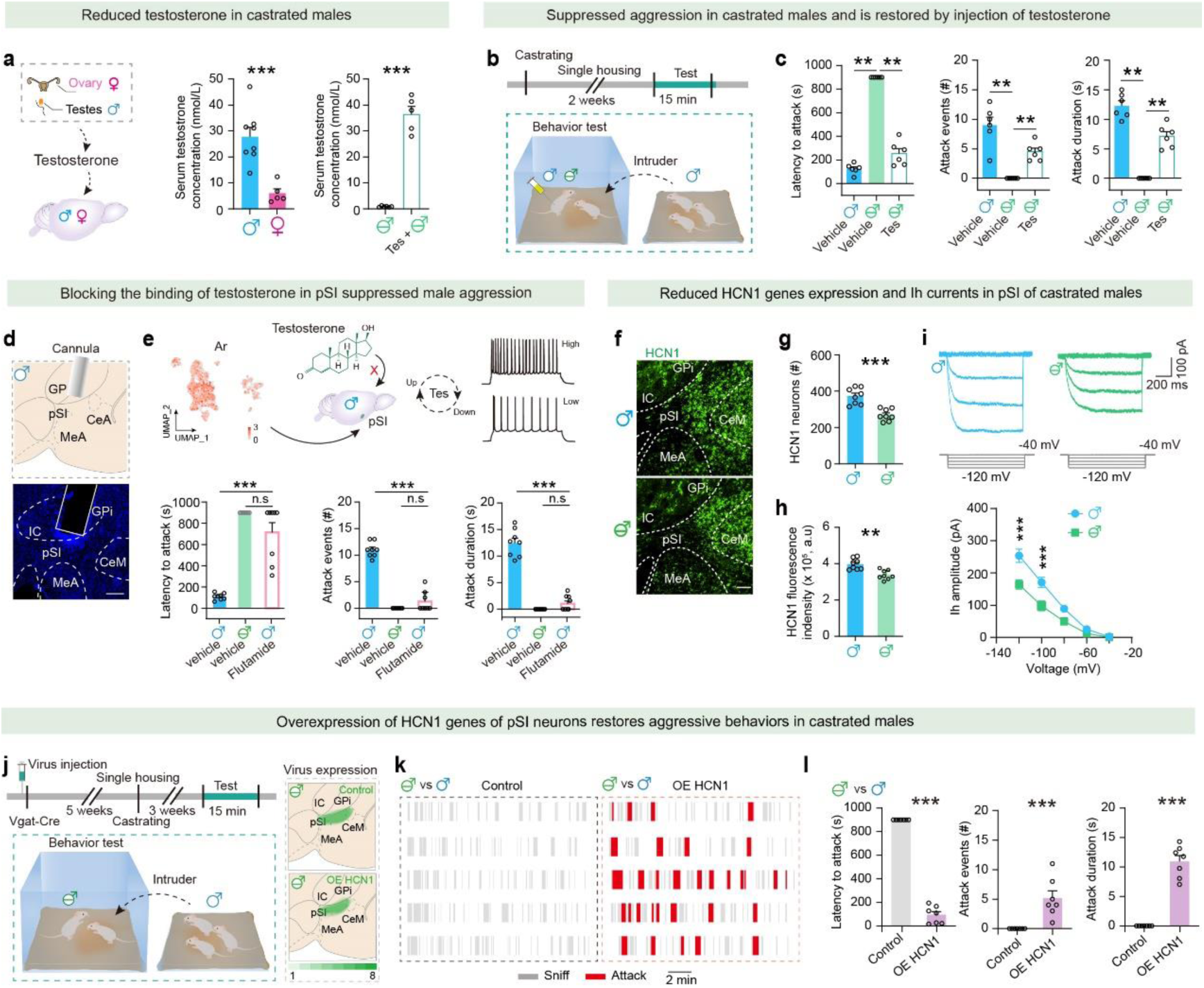
Testosterone facilitates male aggressive behaviors via HCN1 genes in the pSI. (**a**) Summarized testosterone levels in males and females, and the quantification of serum testosterone levels in males, females, castrated males, and castrated males with the injected testosterone. (**b**) Upper, timeline of the animal model of castration for a vehicle or testosterone in a daily injection (100 μg/kg per day, intraperitoneal (i.p.)). Lower, the behavioral paradigm of attack test for singly housed male mice toward a socially housed male intruder in its home cage. (**c**) Effects of intraperitoneally injected testosterone on the first attack latency, attack events, and attack duration of the males administered intraperitoneally with the vehicle, the castrated males administered intraperitoneally with the vehicle, and the castrated males administered intraperitoneally with testosterone. (**d**) Histology, example trajectory trace for inhibiting AR receptor with pharmacological inhibition of AR receptor in the pSI by vehicle control or flutamide. Scale bar, 500 µm **(e)** Effects of pharmacological inhibition of AR receptor in the pSI on the latency to the first attack, attack events, and attack duration of male mice (n=8 mice in each group). (**f**) Representative pSI FISH images showing the HCN1 positive cells (green) in the pSI of male (upper) and castrated male (lower) mice. Scale bar, 100 µm. (**g-h**) Total HCN1 positive cells (f), and fluorescence intensity of positive cells (green) in the pSI in male (n = 3) and castrated male (n = 3) mice (g). (**i**) Representative traces of Ih currents were recorded in male (n = 3 mice) and castrated male (n = 3 mice) mice. Statistical data of the current amplitude (pA) of Ih is plotted against the voltage step in pSI from the male (n = 3 mice) and castrated male (n = 3 mice) mice (n = 18 neurons from 3 mice in each group). (**j**) Behavioral paradigm of attack test for singly housed castrated male mice toward a socially housed male intruder in its home cage after 8 weeks of overexpression HCN1. (**k**) Raster plots of attack behavior recorded during 15-minute encounters in the control and overexpression of HCN1 groups (red blocks indicate the duration of episodes of attack, grey blocks indicate the duration of episodes of sniff). (**l**) Latency to the first attack, attack events, and attack duration at 8 weeks after injection of virus in the control and overexpression of HCN1 groups. Data are mean ± SEM; (a), (c), (e), (g), (h), and (l): Mann-Whitney U test; (i): Two-way ANOVA followed by post-hoc Šidák’s multiple comparisons; *P < 0.05, **P < 0.01, ***P < 0.001, n.s, not significant. For all panels, data are shown as the mean ± SEM. Statistical details are available in the supplementary table S1.

Castration and exogenous testosterone injections are classic methods for altering testosterone levels (Fig. 6a) and affect aggressive behaviors. To test this hypothesis, we performed castration and exogenous testosterone injections and tested the aggressive behaviors in male mice (Fig. 6b). With testosterone levels significantly diminished, we confirmed that castration dramatically reduced aggression (Fig. 6c). Exogenous administered (intraperitoneally) testosterone (100 μg/kg) is effective in regaining high testosterone levels in the castrated males (Fig. 6a). Interestingly, exogenous testosterone administration but not the vehicle injection significantly enhanced castrated male aggression (Fig. 6c), demonstrating that testosterone strongly promotes aggression in castrated males and males. To investigate whether pSI neurons could mediate the testosterone-mediated aggression, we then delivered flutamide, an androgen receptor (AR) blocker into the pSI and tested the aggressive behaviors in the male mice (Fig. 6d). As a result, flutamide (200 μM) reduced male aggressive behaviors through blocking the pSI neurons AR in male mice (Fig. 6e). Therefore, site-specific manipulations of the androgen receptor suggest that the pSI is a critical site through which testosterone exerts its effect in maintaining male aggression.

The essential role of HCN1 in aggression and its sexually differential expression in males and females imply the potential role of HCN1 of the pSI in the hormonal control of aggression. To test this, we then investigated how hormones affect HCN1-mediated male-biased aggressions by examining HCN1 expression when sex hormone levels are altered (Fig. 6f). After the males had been castrated, we found the downregulation of pre-HCN1 mRNA expression of pSI neurons and reduction in Ih currents in males (Fig. 6g-i). Additionally, we performed the slice electrophysiology in the pSI^-PAG^ neurons after injecting retrobeads into the vl/lPAG in sham and castrate males. We recorded pSI^-PAG^ neurons in both voltage-clamp and current-clamp modes and observed significant differences in intrinsic membrane properties of pSI^-PAG^ neurons in sham and castrate males (Extended Data Fig. 10a-k). Specifically, we found that pSI^-PAG^ neurons from castrate males had a population of voltage sag that was approximately 45%, whereas, in uncastrated males, the voltage sag population was around 72.73%. (Extended Data Fig. 10b). Furthermore, we found a significant difference in passive membrane properties between HCN and non-HCN neurons in males and castrated male mice, with the membrane resistances (Rm), membrane time constant (Tau), and membrane capacitance (Cm) being larger in non-HCN neurons than HCN neurons, while the resting membrane potential (RMP) of HCN neurons was more depolarized than non-HCN neurons (Extended Data Fig. 10c-f). By contrast, the electrophysiological properties related to the action potentials (APs) were mostly comparable in HCN and non-HCN neurons in males and castrated male mice. As expected, the minimum current injection needed to produce an AP, known as the rheobase, was higher in castrated males (Extended Data Fig. 10h), indicating HCN neurons in males fire action potentials (APs) after smaller depolarizations. Besides, HCN neurons also exhibited shorter-latency APs and larger half-width APs in males than those of castrated males (Extended Data Fig. 10j-k). Furthermore, to determine how the HCN1 in pSI GABAergic neurons causally affects the hormonal control of aggression, we overexpressed HCN1 in pSI GABAergic neurons by infusing the rAAV-EF1α-Dio-3XFlag-HCN1 or the control rAAV-EF1α-Dio-3XFlag virus bilaterally into the pSI of Vgat-cre castrated male mice and examined the aggressive behaviors 8 weeks after the virus injection (Fig. 6j). Overexpression of HCN1 genes in pSI GABAergic neurons, but not the mice that receive the control virus, was found to be sufficient to enhance castrated male aggression (Fig. 6k-l). Together, these findings suggest that hormonal control of aggression can be mediated by the regulation of the HCN1 gene of GABAergic pSI neurons (Extended Data Fig. 10l).

## Discussion

Our studies have uncovered a molecular substrate within the posterior substantia innominata (pSI) that governs sex-dimorphic aggressive behaviors (Extended Data Fig. 10). A key finding is the higher expression of the HCN1 gene in male GABAergic pSI neurons, linked to increased excitability and contributing to the naturally higher aggression levels in males. Manipulating the HCN1 function allowed us to modulate aggression, revealing its central role in sexually dimorphic behaviors. Silencing HCN1 reduced male aggression, while its overexpression increased aggression in females, demonstrating the gene’s critical role across the sexes. Additionally, testosterone was found to enhance aggression by upregulating HCN1 and remodeling pSI circuits, highlighting the hormone’s role in aggression. Overall, our findings provide a comprehensive understanding of the molecular factors contributing to sex differences in aggression.

### Excitability of an amygdalar circuit regulates sexually dimorphic aggression

Understanding sexual dimorphism in aggression requires examining how aggressiveness is similarly or differently encoded in the neural circuits of males and females (*33–35*). The initiation of aggressive behaviors, which often involve different motor patterns and vary in intensity across sexes, is predominantly governed by shared neural circuits in the brain. It is presumed that sexually dimorphic aggressive behaviors stem from sex differences within these aggression circuits (*2*). Almost all brain regions linked to male aggression also influence female aggression, indicating that the aggression circuitry in rodents is likely qualitatively similar between sexes (*2*). For instance, progesterone receptor (PR) and estrogen receptor type 1 (Esr1)-expressing glutamatergic neurons in the VMHvl are known to regulate inter-male aggression in males and maternal aggression in females (*6, 15, 36*). However, there is a gap in our understanding of how aggression-associated brain circuits trigger differing aggressive behaviors in males and females. Recent findings suggest that these social-related circuits regulate social behaviors by influencing distinct levels of excitability. For example, BNST neurons exhibit higher excitability during mating (*23*), while hyperactivated mPOA neurons correlate more with parenting in females (*37*). In this study, our electrophysiological recordings revealed that male pSI neurons exhibit higher levels of excitability, characterized by more depolarized RMP, increased spiking, and a small rheobase (Fig. 1n-r and Fig. 2k-o). In vivo results further indicate that GABAergic pSI neuronal activity during aggressive behaviors is higher in males than in females (Fig. 1i-m). Additionally, by using optogenetics to activate pSI GABAergic neurons with lasers of varying intensities (Extended Data Fig. 3d-f), we demonstrated that the excitability of pSI GABAergic neurons encodes the intensity of aggression in both male and female mice. As a key node in the circuits regulating aggressive behavior, early circuit mapping has shown that the VMHvl is one of the upstream regions of the pSI. Manipulating the VMHvl-pSI circuit has been found to drive male-biased aggression, with a higher E/I ratio that activates pSI neuronal activity. In contrast, an increased E/I ratio is still insufficient to elicit aggressive behavior in females (*7*), suggesting that there may be other co-factors involved in coordinating neuronal activity, such as the intrinsic electrophysiological properties of pSI neurons found in the present study. Therefore, in addition to the differences in the E/I ratio of integrated synaptic inputs within the pSI, the disparities in the intrinsic electrophysiological properties of pSI neurons further amplify the sexual dimorphism in the aggression circuit regulated by the pSI. Together, our findings of higher intrinsic excitability in male pSI^-PAG^ neurons compared to females suggest this difference in the pSI could account for why natural males are more aggressive than female mice.

### HCN1 channels serve as a molecular substrate in governing sexually dimorphic aggression

Abundant molecular genes such as hormonal signaling, receptors, and ion channels could be involved in sexually dimorphic social behaviors. Current evidence from the ventral medial hypothalamus, ventrolateral part (VMHvl) (*6, 38*), preoptic region (POA) (*39*), and medial amygdala (MeA) (*40*), has revealed significant differences in molecular expression between sexes. Concurrently, increasing evidence shows that specific genes can causally modulate social behaviors. For example, aggressive behaviors can be influenced by hormonal signaling (*1, 6, 11–14*), and primarily through neurons that express hormonal receptors in the brain. Additionally, the role of ion channel expression that mediates a change in ionic conductance of neurons in social neural circuits influencing social behaviors is significant (*2*). Channels such as Cnga (cyclic nucleotide-gated channel) (*25*), or Trpc2 (transient receptor potential cation channel, subfamily C, member 2) (*24, 26*) channels have been identified as crucial in social behaviors, including aggression, and mating behaviors. Additionally, the expression of the hyperpolarization-activated cyclic nucleotide-gated 1 channel (HCN1) gene in the bed nucleus of the stria terminalis (BNST), or lateral septum (LS) influences neuronal excitability and, consequently, affects mating behaviors (*23*) and reward behaviors (*41*).

To gain insight into the molecular mechanisms that underpin the phenotypes outlined above, we focused on the ion channels that govern the intrinsic excitability of neurons. Our study highlights significant differences in biophysical properties such as Rm, RMP, rheobase, and threshold of action potential between HCN and non-HCN pSI^-PAG^ neurons. These findings suggest that the biophysical characteristics of aggression-eliciting neurons in the pSI are associated with HCN channels. RNA-seq analysis revealed that HCN1 is the primary isoform expressed in GABAergic pSI neurons, with HCN2, 3, and 4 expressed at lower levels (Fig. 2f). The increased expression of Ih currents, mediated by HCN channels, plays a key role in governing neuronal excitability. This includes regulating neuronal firing patterns, normalizing temporal summation of synaptic inputs, and facilitating information propagation by integrating calcium signaling. Therefore, the upregulation of HCN channels, particularly HCN1, plays a crucial role in enhancing the excitability of pSI neurons. This heightened excitability is essential for the rapid and efficient propagation of electrical signals within these neurons. Significantly, this mechanism underscores the vital role of HCN channels in facilitating the onset of aggressive behaviors.

HCN channels are involved in a variety of neural functions, including learning, memory, stress, and pain in humans and mice (*20, 21, 42–44*). Abnormalities in these channels are implicated in various neurological disorders, such as epilepsy, depression, and autism (*45–49*). Consequently, the increased expression of Ih currents mediated by HCN channels could play a role in regulating sexually dimorphic aggression. Here we observed impaired HCN1-mediated Ih currents and lower HCN1 channel density in female pSI^Vgat^ neurons (Fig. 3c-f), while castrated males exhibited a significant reduction in HCN1 gene expression (Fig. 6f-i), suggesting a sex-biased correlation in HCN1 expression between males and females. Using CRISPR/Cas9-mediated selective disruption of HCN1 genes in male GABAergic pSI neurons, we achieved a hyperpolarization of these neurons and a reduction in male aggression, similar to the effects observed in females with impaired HCN1 genes. Conversely, overexpression of HCN1 in GABAergic pSI neurons reversed aggression deficits in females and castrated males (Fig. 5). Additionally, the local administration of HCN1 blockers reduced aggression, while the amplification of Ih currents by a cAMP activator increased aggressiveness in males (Fig. 4). These results strongly suggest that aggression circuit signaling converges on the HCN1 channel in GABAergic pSI neurons in both males and females.

Recent advancements in understanding the cellular and molecular mechanisms regulating neuronal excitability have led to the development of small-molecule compounds targeting key regulators like ion channels and receptors (*50*). Our results, which indicate a substantial reduction in male aggression following systemic administration of the HCN blocker ZD7288 (Fig. 4j), underscore the potential translational significance of targeting HCN channels as a strategy to alleviate maladaptive aggression. Notably, substances that inhibit HCN1 channels demonstrate efficient penetration of the blood-brain barrier, facilitating systemic administration in animal models and potentially serving as therapeutic interventions to curb maladaptive aggression.

### Hormonal regulation of sexually dimorphic aggression through HCN1 channels

Hormonal regulation plays a vital role in shaping social behavior, with a well-documented influence on aggression circuits (*12, 16, 32, 51, 52*). Estrogen establishes sexual and territorial behaviors, while testosterone modulates the extent of aggressive displays in males (*12, 13, 32*). Thereby, understanding the integration of molecules with hormonal signals in key social circuits is crucial for comprehending how aggression is modulated. Recent research indicates that sex hormones may modify the excitability of specific brain regions involved in aggression. For instance, testosterone has been observed to increase neural excitability and plasticity in hypothalamic circuits, thereby enhancing aggression (*51*). However, the detailed molecular events downstream of hormone signaling and their interplay with neural circuits that contribute to aggressive behaviors remain largely unexplored (*12, 17, 53*). In our study, we discovered that the depletion of gonadal steroid hormones, specifically testosterone, in adult males markedly reduced male aggressiveness (Fig. 6c) while concurrently decreasing neuronal excitability in pSI^-PAG^ neurons (Extended Data Fig. 10). A notable decrease in aggression was also observed following the local administration of an androgen receptor blocker in the pSI (Fig. 6e), which inhibits the binding of steroid hormones. This finding suggests that hormone receptor signaling within the pSI circuit directly influences aggression. Notably, reduced expression of the HCN1 gene in GABAergic pSI neurons was observed in females and castrated males, indicating that steroid hormones may regulate aggression by affecting the HCN1 channel. The connecting mechanism of the diminished HCN1 expression in the pSI and altered hormone level of two sexes may lie in the hormone receptor neurons, which remains an interesting topic for further investigation.

Collectively, we discovered that GABAergic pSI neurons excitability is a crucial factor in maintaining adult mice in a heightened state of aggression. Elevated expression of HCN1 are effective in regulating cellular firing patterns and excitability. Our findings suggest that the alteration of HCN channels in the pSI circuitry may mediate the sex differences observed in hormone-controlled aggression.

## Acknowledgments

We thank Shari Wiseman for the discussions of the manuscript. We thank Feng Han, Yingmei Lu, Jiadong Chen, Qianqian Ge, and Chong Liu for their technical support. We also thank the Core Facilities of Zhejiang University Institute of Neuroscience for technical assistance. This work was supported by grants from STI2030-Major Projects (2021ZD0203400 to Y.Q.Y.), National Natural Science Foundation of China Major Project (T2293733, T2293730 to Y.Q.Y.), National Natural Science Foundation of China Project (82288101, 82090033, U20A6005 to S.D.), Natural Science Foundation of Zhejiang Province (LZ22H090001 to Y.Q.Y.), Key R&D Program of Zhejiang Province (2024SSYS0019, 2022C03034 to Y.Q.Y.; 2020C03009 to S.D.), CAMS Innovation Fund for Medical Sciences (2019-I2M-5-057 to S.D.), Non-profit Central Research Institute Fund of Chinese Academy of Medical Sciences (2023-PT310-01 to Y.Q.Y.), and Key R&D Program of Guangdong Province (2019B030335001 to Y.Q.Y.).

## Author contributions

Author contributions: K.Y.L. conducted in vitro electrophysiological recording and single nucleus sequencing experiments and analyzed data; K.Y.L., Q.W., L.N.P., J.R.W., L.M., Z.G.Z., and H.F.L. performed the immunostaining experiment and behavior tests; G.C.D. performed the RT-qPCR; K.Y.L., Z.G.Z., and Q.W. performed the analysis of data. X.M.L., L.S., Y.J.L., and H.L.Z. contributed to the discussion of the results. K.Y.L., Z.G.Z., S.D., and Y.Q.Y. designed and wrote the manuscript.

## Data availability statement

All datasets used in this study are in the process of being deposited to public repositories. All script files used in the analysis in this manuscript will be available upon submission.

## Materials and Methods

### Animals

Mice C57BL/6J, NIH (S), Vgat-Cre mice, Vgat-Venus, and Rosa26-LSL-Cas9 knock-in mice (JAX#024857) of both sexes were used in the present study. For behavior tests in mice and functional manipulation using in vivo fiber photometry and in vivo recording experiments, mice 2-4 months old were used. Mice were housed under a 12-hour light-dark cycle (lights on from 7 am to 7 pm), at a room temperature of 22±1 °C and a humidity level of 55±5%. Food and water were available and ad libitum before behavioral training. Animal experiments were conducted in accordance with the Guidelines for the Care and Use of Laboratory Animals of Zhejiang University.

### Viruses

AAV_2/9_-hSyn-Flex-ChrimsonR-tdTomato-WPRE (1 × 10^12^ genomic copies/ml), AAV_2/9_-hSyn-DIO-hM3Dq-mCherry (2.0 × 10^12^ genomic copies per ml), AAV_2/9_-CAG-DIO-GCaMP6m-WPRE (1.0 × 10^13^ genomic copies/ml), and AAV_2/9_-Syn-Cre-EGFP (1.0 × 10^13^ genomic copies/ml) were made by Taitool Bioscience Co., Ltd. (Shanghai, China). rAAV_2/9_-U6-spsgRNA (HCN1)-U6-spsgRNA(HCN1)-CMV-EGFP-bGH (2.0 × 10^12^ genomic copies/ml), rAAV_2/9_-U6-spsgRNA(scramble)-U6-spsgRNA(scramble)-CMV-EGFP-bGH (2.0 × 10^12^ genomic copies/ml), were provided by BrainVTA Co., Ltd. (Wuhan, China). rAAV_2/9_-U6-EF1α-Dio-3Xflag-HCN1-WPREs (5.0 × 10^12^ genomic copies/ml), rAAV_2/9_-U6-EF1α-Dio-EGFP-WPRE-hGH (5.0 × 10^12^ genomic copies/ml), were provided by BrainVTA Co., Ltd. (Wuhan, China).

### Stereotaxic surgery

Mice were anesthetized with isoflurane (1-2%) during stereotaxic injection. Depending on the viral titer and the desired expression strength, we injected 80-150 nl of virus solution into each targeted location at a rate of 20 nl/min. After the injection, the syringe was kept in place for 5-10 minutes before being removed, and a heating pad was used to maintain the core body temperature at 36 °C. The coordinates of viral injection sites including the PAG (AP, -4.70 mm; ML, ±0.8 mm; DV, -2.8 mm), pSI (AP, -0.90 mm; ML, ±2.3 mm; DV, -4.5 mm). The coordinates of the optical fiber placement sites in PAG or pSI was placed at 200 μm above the virus injection sites. A cannula was implanted in pSI (AP, -0.90 mm; ML, ±3.1 mm; DV, -3.8 mm) at a 10° angle for in vivo drug administration. All animal controls in this study were animals with the same genetic background injected with control viruses. All mice returned to their home cages after surgery. Mice were allowed to recover for at least 3 weeks before being used for behavioral studies.

### Histology and immunohistochemistry

All mice were histologically verified for viral expression and implant placement after completing the behavioral experiments. Only mice with the confirmed correct viral expression and fiber implant placement were used for analysis.

### Tissue preparation

After experiments, mice were perfused with saline followed by 4% paraformaldehyde in 1×PBS. The brain was then removed and placed in 4% PBS at 4 °C for overnight fixation. Following fixation, the brain was cryoprotected in 15% to 30% sucrose (wt/vol) at 4 °C. Free-floating coronal sections (35 μm) were cut by a microtome (Thermo Fisher Scientific, Waltham, MA) for c-Fos immunohistochemistry or FISH.

### c-Fos immunohistochemistry

To induce aggression in mice for c-Fos quantification of both sexes, we introduced a socially adult male intruder (C57BL/6N, 6-8 weeks) into the home cage of a sexually experienced adult mouse (C57BL/6N, 10-12 weeks). Mice were exposed to aggression for 10-15 min. In the control group, an intruder male was introduced to be investigated by the male subject mice but with no attack observed. These mice were perfused 1.5 h later, and brain sections were cut.

After rinsing with 0.3% Triton-X 100 (vol/vol) in 1×PBS (30 min) and blocking with 10% (wt/vol) normal bovine serum for 1 h at room temperature, sections were incubated with the following primary antibodies (12-24 h at 4 °C), anti-c-Fos (1:1500, rabbit, Calbiochem). After rinsing, sections were incubated with fluorophore-conjugated secondary antibody (1:2000; Millipore) for 2 h at room temperature. Antibodies were diluted in 1×PBS containing 5% BSA and 0.2% Triton X-100. After the anti-c-Fos immunohistochemistry reaction, nuclei were stained with DAPI, and confocal images were captured using a 10×, 20× objective (Olympus FV-1200), cell counting was carried out manually using Fiji (NIH). For each mouse, we quantified 3 sections from RI-tested mice that specifically covered the pSI area and quantified the numbers of anti-c-Fos cells with the boundary of the pSI using Fiji (NIH). Statistical significance was assessed using the Mann-Whitney U test.

### Fluorescent in Situ Hybridization

Fluorescence in situ hybridization (FISH) was performed to examine the expressions of c-Fos, Slc32a1, Slc17a6, and HCN1 in the pSI. All reagents and probes are commercially available from Advanced Cell Diagnostics. The RNAscope Fluorescent Assay is a FISH technique used to visualize cellular RNA targets in frozen tissue. RNAscope in situ hybridization was performed by collecting frozen coronal sections (35 μm) as described above. Hybridizations were performed using RNAscope Multiplex Fluorescent Assay (ACDBio) according to the manufacturer’s instructions. ACDBio probes used were Cfos (#316921-C3), Slc32a1 (#319191-C1), Slc17a6 (#319171-C3) and HCN1 (#423651-C2). Probes were visualized with ACDBio TSA fluorescent dyes. Finally, the glass coverslips were mounted onto glass slides. Images were acquired using a 10×, 20× objective (Olympus FV-1200). For each group, we quantified 3 sections from 3-5 mice that specifically covered the pSI area and quantified the cell numbers of HCN1-, Vgat-, and Vglut2-positive cells with the boundary of the pSI using Fiji (NIH).

### Slice Electrophysiology

Mice (8-10 weeks) were deeply anesthetized with isoflurane and and perfused with ice-cold slicing solution via heart. The whole brain was quickly dissected into ice-cold cutting solution containing the following (in mM): 93 N-methyl-D-glucamine, 2.5 KCl, 1.2 NaH_2_PO_4_, 20 HEPES, 25 D-glucose, 30 NaHCO_3_, 10 MgSO_4_, 0.5 CaCl_2_, 5 sodium ascorbate, 3 sodium pyruvate, and 1 kynurenic acid (pH: 7.3-7.4; mOsm: ∼300). Then the brain was cut coronally into 300 μm slices on a microtome (VTA-1200S; Leica). Slices containing the pSI were incubated at 33 °C in this solution for 15 min, then transferred to a similar solution containing (in mM): 93 NaCl, 2.5 KCl, 1.2 NaH_2_PO_4_, 20 HEPES, 25 D-glucose, 30 NaHCO_3_, 2 MgSO_4_, 2 CaCl_2_, 5 sodium ascorbate, 3 sodium pyruvate, and 1 kynurenic acid (pH: 7.3-7.4; mOsm: ∼300), and incubated for at least 1 h at room temperature (RT) (24-26 °C). Experiments were performed in ACSF containing (in mM): 125 NaCl, 3 KCl, 1.25 NaH2PO4, HEPES, 10 D-glucose, 26 NaHCO_3_, 2MgSO_4_ and 2 CaCl_2_, heated to 33 °C. All the solutions were gassed with 95% O_2_ and 5% CO_2_. The slices were transferred to a recording chamber on the stage of a fluorescence microscope (BX51WI; Olympus). Patch electrodes were pulled from borosilicate capillaries (BF150-86-10; Sutter instrument) and had impedances of 3-5 MΩ when filled with intracellular solution containing (in mM): 135 K-gluconate, 6 NaCl, 10 HEPES, 0.5 EGTA, 10 Na_2_-phosphocreatine, 4 Mg-ATP, 0.3 Na_2_-GTP (pH:7.2 adjusted with KOH; mOsm: 290-300) for recording pSI^-PAG^ neuronal action potentials. Recordings were made with a MultiClamp 700B amplifier (Molecular Devices) and Digi-data 1440A interface (Molecular Devices). Signals were low pass filtered at 10 kHz and digitized at 10 kHz (MICRO3 1401, Cambridge Electronic Design). Series resistance did not exceed 25 MΩ, and input resistance was more than 300 MΩ. The data were acquired and analyzed using Clampfit 10.6.2.2 software (Cambridge Electronic Design).

To characterize the spiking patterns of pSI^-PAG^ in females and males, we first injected red retrobeads into vl/lPAG and performed whole-cell patch clamp recordings on red retrobeads-expressing neurons in the pSI of the wild-type males (n = 10 mice) and females (n = 10 mice). To test the hypothesis that the differences in electrophysiological properties related to action potential (AP). We extracted electrophysiological features related to passive membrane properties, firing patterns, and action potential (AP) were extracted. All the properties of action potential (AP) in the present study were determined by analyzing the first spike elicited by the injection of the depolarization current step. The temporal sequence of AP or spiking pattern. Graphs of the membrane potential slope (dV/dt) between male and female pSI^-PAG^ neurons. The spike latency and half-width of AP, the intrinsic biophysical properties of a neuron, such as resting membrane potential (REM), membrane resistance (Rm), membrane time constant (τm), membrane capacitance (Cm) of pSI-PAG neurons, the voltage threshold for action potential (Vth), and the minimum injected current that needed to produce an action potential (rheobase), were analyzed.

To activate ChrimsonR-expressing neurons, 5-ms pulses of full-field illumination (5 mW) were delivered onto the recorded cell with a red LED light at different intervals (1s, 100 ms, 50 ms, 25 ms). For optogenetically-evoked synaptic currents recordings, pipettes were filled with a solution made up of the following components (in mM) 135 CsMeSO_3_, 10 HEPS, 0.5 EGTA, 3.3 QX-314, 4 Mg-ATP, 0.3 Na_2_-GTP, 8 Na_2_-Phosphocreatine (pH:7.2 adjusted with CsOH; mOsm: 290-300). To activate ChrimsonR-expression axons, 5-ms pulses of full-field illumination (5 mW) were delivered onto the recorded cell with a red LED light. oEPSC and oIPSC were recorded by holding the membrane potential of recorded neurons at − 70 mV and 0 mV, respectively. To examine the monosynaptic nature of oIPSC in vlPAG following light activation of pSI inputs, slices were perfused with TTX (1 μM) to block the sodium channel and TTX followed by the addition of 4-aminopyridine (4-AP, 100 μM), a potassium channel blocker, to facilitate GABA release from synaptic vesicles. TTX was purchased from Tocris Bioscience (Cat. no. Tocris_1078). 4-AP was purchased from Sigma-Aldrich (Cat. no. 275875). Bicuculline (100 μM) was bath-applied to block GABA_A_ receptors. HCN neurons and non-HCN neurons were identified by voltage sag and rebound depolarization mediated by the Ih currents. Brief (1 s) negative current pulses revealed a depolarizing “sag” (indicative of the hyperpolarization-activated cation current, Ih), followed by a post inhibitory rebound. The size of the voltage sag in response to hyperpolarizing direct-current steps of 1 s was calculated as the amplitude of the voltage sag (mV). In response to a current injection of -100 pA, neurons with a peak voltage deflection divided by the amplitude of the voltage sag deflection greater than 10% were considered to express substantial Ih current. In Figure. 3a, the HCN channel blocker ZD7288, was applied to pSI^-PAG^ neurons. The effect of ZD7288 was determined by comparing the averaged Ih currents over a 3 min period just before ZD7288 application with averaged Ih currents during the 5-10 min period after ZD7288 application.

### Analysis of Electrophysiological Parameters

All electrophysiological data were analyzed with ClampFit 10 (Molecular Devices) and Mini analysis 6.0 (Synaposoft). Projected positions of these neurons in a reference brain atlas show that they are tightly clustered within the pSI as defined by the atlas and there is no systematic deviation between the two sexes. The resting membrane potential (REM) was measured after the formation of the whole-cell patch clamp, without current injection. Input resistance (Rm) was calculated using linear regression established between electronic voltage response to a hyperpolarization currents step (from 0 pA to − 30 pA in − 5 pA increments, 1 s). The membrane time constant (τm) was obtained by averaging the voltage response to a hyperpolarization current step (− 20 pA, 500 ms, ∼10 repeats). Specifically, Rm was determined by the voltage response at the steady state to the current while the τm was determined by fitting this average response to a single exponential curve. The membrane capacitance (Cm) was calculated according to the formula Cm = τm/Rm. All properties of action potential (AP) were determined by analyzing only the first spike elicited by the injection of the depolarization current step. Rheobase was the minimum current that elicited a spike when holding the membrane potential at resting membrane potential. Spike threshold (Vth) was defined as the voltage when dV/dt equals 20 V/s. The spike peak was the maximum voltage during the AP, and spike amplitude was measured by subtracting the Vth from the spike peak. Spike latency was computed as the time from the onset of a current pulse to the peak of the AP. Spike half-width duration was defined as the width at half-spike amplitude above the threshold. The relationship between the injection current and action potential firing frequency (I-F curve) was measured from the first trace injection current 0 to the last trace before a prominent reduction of the AP firing. Whole-cell recordings were rejected if the initial membrane potential was more negative than − 80 mV or if membrane potential changed by >10% and input resistance changed by 30% during the recording.

### Monosynaptic retrobeads tracing

Monosynaptic retrobeads tracing experiments were performed in adult wild-type mice. Mice were anesthetized and injected, as described above, with 80 nl red retrobeads (Lumofluor, USA) into predicted pSI output sites: vl/lPAG (coordinates: AP: − 4.7, ML: ±0.8, DV: − 2.8). Two weeks after the injection, mice were used for patch clamp recording.

### Behavioral tests and data analysis

#### Fiber photometry signal acquisition and analysis

Following AAV-EF1a-DIO-GCaMP6m injection in Vgat-Cre male and female mice, an optical fiber (2.30 μm OD, 0.37 NA) was placed in a ceramic ferrule (2.5 mm OD, 1.26 mm ID) and inserted toward the pSI at 200 μm above the virus injection sites. A skull-penetrating M1 screw and dental acrylic were used to secure the ferrule in place. To facilitate recovery and AAV expression, mice were housed individually for 2 weeks after virus injection. A fiber photometry system (Thinker Tech Nanjing Biotech Ltd, Nanjing, China) was used for recording. To record GCaMP6m fluorescence signals, the beam from a 488 nm laser (OBIS 488LS; Coherent, Santa Clara, CA, USA) was reflected with a dichroic mirror and focused with a 10× objective lens (NA = 0.3; Olympus, Tokyo, Japan). An optical fiber (2.30 mm OD, NA = 0.37; 2 m long) guided the light between the commutator and the implanted optical fiber. The laser power at the tip of the optical fiber was adjusted to a low level (0.03-0.05 mW) to reduce GCaMP bleaching. The GCaMP6m fluorescence signal was filtered with an EYFP bandpass filter and collected by a photomultiplier tube (R3896; Hamamatsu Photonics). An amplifier converted the photomultiplier current output to a voltage signal, which was further filtered through a low-pass filter (40 Hz cut-off; Brownlee 440). The analog voltage signals were digitized at 500 Hz (Power 1401 digitizer, Cambridge Electronic Design) and sampled with software (TDMS). Data from fiber photometric recording were exported as MATLAB files for further analysis. Fiber photometry-related aggression behavioral data were analyzed using MATLAB. After being segmented and aligned to the onset of behavioral events within individual trials or bouts, all of the raw fluorescence data were smoothed with a moving average filter (10 ms span). The fluorescence change values (△F/F) were calculated as (F–F0)/(F0– V_offset_), where F0 is the baseline fluorescence signal averaged over a 2 s/4 s time-window before a trigger event and V_offset_ is the fluorescence signal recorded before the cannula was connected to the optical fiber above the pSI. △F/F values are presented as heat maps or average plots with a shaded area indicating the SEM.

During the test, the aversive stimuli were air puffs (delivered by a hand-pumped compressor with its opening 3 cm from the mouse’s nares; one press of the pump generated a gentle 0.5 s puff) and a flying object (a flattened glove or cotton ball was introduced into the tested mouse’s cage above its head, and then moved towards its body) for Vgat-Cre males and females. The onset of each stimulus triggered data alignment. Social or non-social entry refers to the threat period after an intruder male or object was first introduced into the home cage of a subject Vgat-Cre mouse. The timing of each entry triggered the alignment of the data. Aggressive social behaviors of these subject Vgat-Cre mice were recorded during the application of various contextual cues.

We calculated the average △F/F during the following stimuli and associated aggressive behaviors: offensive inter-male attack, female-male attack, sniff without attack (a sniff not followed by an attack within 5 s), threat (subject mouse threatened other males with boxing postures but no bite), defensive attack, and multiple social and non-social cues based on the peristimulus time histograms (PSTHs). The non-parametric Wilcoxon signed-rank test or t-test was used to determine whether the area under the curve (AUC) per second and peak △F/F before, during, and after events were significantly different between males and females.

#### ChrimsonR-mediated neural activation

To acutely manipulate the GABAergic pSI neurons, Vgat-Cre mice of both sexes unilaterally stereotaxically injected AAV_2/9_-hSyn-Flex-ChrimsonR-tdTomato-WPRE virus (80 nl/side) into the pSI, at 8-10 weeks, and the viruses were allowed to incubate for 3 weeks before behavioral testing or perfusion while the animal recovered in its home cage. After the recovery period, animals were tested. A ferrule patch cord was coupled to the ferrule fiber implanted in the mouse using a zirconia split sleeve (Doric Lenses). Optic fibers were connected using an FC/PC adaptor (Doric Lenses) to a 632-nm red laser (Inper Laser). For stimulation experiments, red (632 nm) light was delivered in 5 ms pulses at 40 Hz for 15 s, with an intensity of 3 mW at the fiber tip for all experiments other than the testing scalability. To further investigate how the pSI ^Vgat^ neurons contribute to aggression, we introduced an adult male intruder into the cage of subject mice and accessed aggression behaviors during photoactivation of the pSI^Vgat^ neurons or pSI^Vgat^-VL/LPAG terminals. Stimulations were delivered based on the animal’s behavior, with at least one minute between each stimulation trial. In particular, the probability of observing stimulation-induced aggression was lower when animals were on the opposite side of the cage or not facing the intruder; therefore, we triggered stimulation during random bouts when the animal was in the proximity of intruders. The same criteria were used for both male and female mice.

Adult males and females were moved to a behavioral testing room in the behavioral testing cage and given more than 1 h to adapt before testing. The cages were placed on a custom behavior rig equipped with video acquisition capabilities before testing began. Following patch cord attachment to the implanted fiber, animals were given more than 5 minutes before the intruder was placed into the behavioral testing cage.

#### HM3Dq-mediated neural activation

To activate the GABAergic pSI neurons, the AAV-hSyn-DIO-hM3Dq-mCherry (a Cre-dependent pharmacogenetic activating virus, 80 nl/side) was directly used to unilaterally infection the pSI neurons in Vgat-Cre mice. To minimize stress with fear or anxiety, mice were handled daily in a manner that simulated injection for at least 7 days before testing. Three weeks after the injection, mice were moved to a behavioral testing room in a behavioral testing cage and given more than 1 h to adapt before testing. The cages were placed on a custom behavior rig equipped with video acquisition capabilities before testing. The following mice were intraperitoneally injected with either saline or CNO (0.5 mg/kg), respectively. Thirty minutes after the intraperitoneal administration of the drug, an adult intruder mouse (20-30 g) was introduced for approximately 15 minutes to evaluate the aggression level of the mice.

#### CRISPR/Cas9-mediated HCN1 gene disruption in GABAergic pSI neurons

The target sites of HCN1 genes for Staphylococcus aureus CRISPR/Cas9 were designed using GHOPCHOP (http://chopchop.cbu.uib.no). the target sequences were as follows: sgHCN1: 5’-CAACTCCGTGTGCTTCAAGG-3’ and sgHCN1: 5’-AGACGCTGTTGCCATCGTCG-3’. Oligonucleotides encoding guide sequences were purchased from Sigma-Aldrich and cloned individually into the BsaI fragment of pX601 (Addgene plasmid 61591). U6-sgHCN1 fragments were PCR-amplified, respectively using pX601-sgHCN1 as a template. Amplified fragments were cloned tandemly into MluI-digested rAAV-CMV-Cre-EGFP by the Gibson assembly method. AAV constructs carrying nontargeting guide sequences (5′-GCGAGGTATTCGGCTCCGCGT-3′) were used as control. Next, the plasmid was digested with MluI and subjected to self-ligation to remove U6 promoter and RNA (sgRNA) scaffold sequences, rAAV-U6-sgHCN1-U6-sgHCN1-CMV-Cre-EGFP (sgHCN1) and rAAV-U6-sgControl-U6-sgControl-CMV-EGFP (sgControl) were packaged into AAV-DJ by BrainVTA Co., Ltd. (Wuhan, China).

Rosa-LSL-Cas9 males (8-10 weeks old, n = 16) were separated into two groups in a random manner (n = 8 in each group). Under anesthetics and analgesic, according to the pSI coordinates as described above, one group received a bilateral stereotaxic injection of 80 nl (each side) sgControl (3.08 × 10^12^ gc/ml) mixture of AAV-hsyn-Cre as the control group. The other group received a bilateral stereotaxic injection of 80 nl sgHCN1 (3.47 × 10^12^ gc/ml) mixture of AAV-hsyn-Cre to monitor the effect of pSI neuron selective HCN1 gene disruption on aggressive behaviors. The experimental group consisted of Vgat-Cre mice that bilaterally expressed Caspase3 and Vgat-Cre::Rosa-LSL-Cas9 mice that bilaterally expressed sgHCN1 in the pSI. The control group consisted of Vgat-Cre mice that bilaterally expressed EGFP and Vgat-Cre::Rosa-LSL-Cas9 mice that bilaterally expressed sgControl in the pSI. The designs of sgHCN1 and sgControl are described in “CRISPR/Cas9-mediated HCN1 disruption in pSI^Vgat^ neurons^”^ below. After surgery, mice were singly housed with food and water *ad libitum* to recover. Resident-intruder testing was recorded continuously from week 3 onwards, with weekly updates up to week 8 (Resident-intruder testing lasted until 12 weeks in half of each group) following surgery. Following recording in 8 to 12 weeks after virus injection, slices were prepared from each group for an in vitro electrophysiology experiment to determine the RMP and firing properties of the GABAergic pSI neurons labeled by the EGFP flag.

#### Overexpression of HCN1 gene in GABAergic pSI neurons

The truncated coding sequence of mice HCN1 (1-2733 bp) was cloned into the PT-4151 vector to generate PT-HCN1-3XFlag. S3-S4 linker residues were then strategically deleted by site-directed mutagenesis to create PT-EF1α-DIO-3XFlag-HCN1-WPREs, allowing the expression of HCN1 that favors channel opening. rAAV-EF1α-DIO-3XFlag-WPREs and rAAV-EF1α-DIO-3XFlag-HCN1-WPREs were generated by co-transfecting HEK-293T/17 cells with pDG9 helper plasmid. The recombinant products were then harvested by multiple freeze-thaw cycles and purified using density gradient ultracentrifugation and dialysis, yielding rAAV of concentrations on the order of 1012 genome copies (gc)/mL. rAAV-EF1α-DIO-3XFlag-WPREs and rAAV-EF1α-DIO-3XFlag-HCN1-WPREs were packaged into AAV-DJ by BrainVTA Co., Ltd. (Wuhan, China).

16 young (8-10 weeks old) Vgat-Cre females were separated into two groups randomly (n=8 in each group). Under anesthetics and analgesic, according to the pSI field coordinates as described above, one group received bilateral stereotaxic injection of 80 nl (each side) rAAV-EF1α-DIO-3XFlag-WPREs (2.5 × 10^12^ gc/ml) as the control group. The other group received a bilateral stereotaxic injection of 80 nl rAAV-EF1α-DIO-3XFlag-HCN1-WPREs (2.5 × 1012 gc/ml) to monitor the effect of HCN1 gene overexpression specifically in pSI Vgat^+^ neurons on aggressive behaviors. After surgery, mice were singly housed with food and water *ad libitum* to recover. Resident-intruder testing was recorded continuously from week 3 onwards, with weekly updates up to week 8 (Resident-intruder testing lasted until 12 weeks in half of each group) following surgery. After the recording at week 8 to 12 weeks post-viral injection, slices were prepared from each group for an in vitro electrophysiology experiment to determine RMP and firing properties of the GABAergic pSI neurons.

#### Cannula implantation and drug infusion

ZD7288 (Cat. no. Z3777), and zatebradine hydrochloride (Cat. no. Z0127, referred to as zatebradine) were purchased from Sigma-Aldrich. Ivabradine hydrochloride (Cat. no. S2086, referred to as ivabradine) and Flutamide (Cat. no. S1908) were purchased from Selleck.cn. The cannula was stereotaxically bilaterally implanted above the pSI. The inner and outer diameters of the cannula were 150 μm and 300 μm, respectively. The cannula was fixed to the skull using acrylic cement. During drug infusion, the cannula was connected with a catheter filled with drugs (ZD7288, 100 μM; zatebradine, 100 μM; ivabradine, 100 μM; Flutamide, 100 μM) or vehicle for injection. The other end of the catheter was connected to a Hamilton syringe controlled by an infusion pump to drive the delivery of the drug or vehicle (50 nl/min). Aggressive behaviors were measured 30 min after infusion was completed (300 nl). Flutamide was injected continuously every day for up to 2 weeks. At the end of the experimental session, the test mice were perfused, and the coronal brain sections containing pSI were inspected for the presence of a cannula track. Mice without a cannula track above the pSI were excluded from further analysis.

50 mM ZD7288 stock solution (dissolved in DMSO), zatebradine, and ivabradine were prepared in saline with a concentration of 0.5 mg/ml for in vivo experiment with a different dosage of 0 mg/kg, 0.2 mg/kg, 1 mg/kg, 2 mg/kg, 5 mg/kg (0.1 ml/10 g, intraperitoneally), respectively. Testosterone Enanthate (Cat. no. S3717, referred to as Testosterone) was purchased from Selleck.cn, was prepared at a concentration of 1mM/L in 2% DMSO + 30% PEG300 + 5% Tween80 +ddH_2_O for *in vivo* experiment with a dosage of 0.05 mg/kg (0.1 ml/10 g, intraperitoneally). Testosterone solution was ultrasonicated before application and injected continuously every day for up to 2 weeks. To minimize stress and fear or anxiety, mice were handled daily in a manner that simulated injection for at least 7 days before testosterone or vehicle treatment.

#### Open field test

Mice were placed in an evenly illuminated 50 cm × 50 cm open-field test box. The mouse was placed in one of the corners of the arena at the start of each session and was allowed to explore the arena for 5 min being recorded using a webcam and Viewer software (Biobs kerve). The arena was cleaned thoroughly between each trial.

#### Behavioral Measurements and Video Analysis

For behavioral analysis, we manually carried out frame-by-frame annotation of aggression and stimulation bouts. To compare and combine data across all trials, trials from all animals were aligned to the onset of stimulation. Each frame during each trial was scored as showing the specified behavior or not. This approach allowed for comparison across animals and trials, and a “percentage of trials showing behavior” could be calculated.

Behavioral parameters such as the possibility of trials of induced attacks, the number of attacks, the latency to the first attack, and the total duration of the attacks were analyzed only in animals that displayed aggressive behaviors. Features of the aggressive behavior paradigm in our experiment include (a) Various social-contextual conditions, including singly housed for 2 weeks or socially housed subject mice before the behavior test. During the test, the mice were introduced to a novel cage or as a resident in their home cage for the aggression test. (b) The tested mice, which included adult males and females, were used as the test subjects. Each test ran for 15 min and was videotaped and scored for the following parameters: mounting and mounting with a pelvic thrust. All videos were recorded at 30 fps, manual annotation was performed by an observer blind to experimental conditions, and annotations were then processed and analyzed using customized MATLAB programs to characterize and quantify behavioral episodes as previously described.

#### Gonadectomies and testosterone treatment

Adult mice were anesthetized with sodium pentobarbital (1%) before the testes were removed. In testosterone supplement studies, testosterone was dissolved in 2% DMSO, 30% PEG300, 5% Tween − 80 and diluted in ddH_2_O (Sigma). Mice received a subcutaneous injection of either 0.5 ug testosterone in 100 μl saline or 100 μl of saline with an equivalent amount of 2% DMSO, 30% PEG300, 5% Tween 80, ddH_2_O.

#### Blood sampling and testosterone ELISA

Whole blood samples of 600-800 μl were collected from the submandibular vein of wild-type mice. Serum testosterone (ENZO, Testosterone Parameter Assay Kit) levels were assessed via ELISA according to manufacturer specifications. Briefly, blood was collected in a serum separator tube, allowed to clot for 2 hours at room temperature, centrifuged at 1,000 g for 15 min, and stored frozen (− 80 °C) until analysis. Hormone titers were assayed with kits according to the manufacturer’s protocols. To assay, the physiological testosterone level associated with the induced aggression in males, females, castrate males, and testosterone-administrated castrate males, and after the experiment, the mice were perfused, and plasma concentrations of testosterone were collected.

#### Quantitative RT-qPCR

Total RNAs from brain tissues was extracted using TRIzol (Takara Bio). RNAs were reverse-transcribed using PrimeScript RT Master Kit (Takara Bio). Real-time PCR was performed using the TB Green Premix Ex Taq (Tli RNaseH Plus) on a Bio-Rad CFX96 Real-Time PCR Detection System (Bio-Rad). Primer sequences were as follows: Forward: AGCCAGAGCACTTCGTATCG, Reverse: CACTGGCGAGGTCATAGGTC. The relative expression of target RNAs was measured using the 2^− ΔΔCt^ method. ΔCt = Ct _(Target genes)_ – Ct_β-actin_. ΔΔCt = ΔCt _(Target genes)_ – ΔCt _(average ΔCt of control)_. Relative fold changes were determined by 2^− ΔΔCt^ and normalized to the expression levels of β-actin.

#### Sample collection and single-nucleus RNA sequencing

All procedures involving C57BL/6 J mice (Jackson Labs, #000664) were conducted in accordance with the ethical guidelines and received approval from the Animal Advisory Committee at Zhejiang University. For single-nucleus RNA sequencing (snRNA-seq), male and female mice, aged eight weeks, were anesthetized deeply using 1% sodium pentobarbital (dose: 0.1 g/kg body weight) and transcardially perfused with ice-cold oxygenated N-methyl-D-glucamine (NMDG)-based artificial cerebrospinal fluid (ACSF). The ACSF solution consisted of 93 mM NMDG, 2.5 mM KCl, 1.25 mM NaH_2_PO_4_, 30 mM NaHCO_3_, 20 mM HEPES, 25 mM glucose, 5 mM sodium ascorbate, 2 mM thiourea, 3 mM sodium pyruvate, 10 mM MgSO_4_.7H_2_O, and 0.5 mM CaCl_2_.2H_2_O. Mice were rapidly decapitated, and whole brains were extracted into ice-cold oxygenated NMDG-based ACSF. Subsequently, 200 μm thick coronal sections were cut in the cold NMDG-based ACSF with a vibratome slicer (VT1200s, Leica Biosystems). Carbogen (95% O_2_, 5% CO_2_) was bubbled continuously during sectioning. The pSI sections were snap-frozen in liquid nitrogen and stored at − 80 °C. Dissected mouse samples were utilized for nuclei isolation. Nuclei were isolated and purified following the 10× Genomics protocols (CG000393 • Rev A), with some modification. The nuclear suspension was diluted to a concentration of 500–1000 nuclei/μL before being loaded onto the 10× Chromium instrument. Nuclear suspension loading onto the Chromium Next GEM Chip, nuclear barcoding, cDNA amplification, and library construction were carried out according to the 10× Genomics protocols (CG000204 Rev D). Libraries were sequenced on the Illumina NovaSeq 6000 System at LC-Bio Technology Co., Ltd. in Hangzhou, China.

#### SnRNA-seq data processing

The snRNA-seq data constructed using 10× Genomics were processed through the Cell Ranger Single-Cell Software Suite (v7.1.0), which performed sequencing read alignment and gene expression quantification. We used the *CellRanger aggr* pipeline to merge cells from the same species under different samples, with the parameter “*normalize* = *none*”. The STAR algorithm was used to align the FASTQ output, which were obtained from sequencing the library constructed from the nuclear suspension, to the GRCh38 reference genome. Next, gene-barcode matrices were generated for each sample by counting the unique molecular identifiers (UMIs), barcode count, and genes without expression across all cells were removed. Finally, we generated a gene-barcode matrix that contained barcoded cells and gene expression counts. All additional analyses were performed using the Seurat (v4.3.1, http://satijalab.org/seurat/) and RTool 4.3, including quality control and all subsequent analyses. To eliminate the influence of low-quality cells, such as empty droplets and multiplets, cells with expressed genes <500 or >8000 were excluded. The percentage of UMIs mapped to mitochondria was set to less than 25%. Finally, a total of 15719 cells after quality control were used for further downstream analysis, including 8488 cells from male samples and 7271 cells from female samples. We identified the top 2000 variable features using the “vst” method for each dataset. Datasets were then anchored and integrated using the integration procedure from the Seurat package to eliminate the batch effects among the samples. ScaleData function was used to perform a linear scaling transformation on the identified variable features using default parameters. Principal component analysis (PCA) was performed on the scaled data to reduce the dimensionality. The statistical significance of the PCA scores was determined using the JackStraw function. The first 25 principal components were used for identifying the neighbors and clustering the cells with a resolution of 0.6. The cell clusters were visualized using 2D uniform manifold approximation projection (UMAP) plots. The FindAllMarkers function was used to identify the genes specifically expressed in each cell cluster. We identified the cell types based on the expression of well-established gene markers. Cells were divided into 21 main cell lineages, of which 8698 were neurons cells. To compare the characteristics of pSI cells from male and female samples, we integrated pSI cells from 20 males and 20 females’ samples and identified 21 clusters by unsupervised clustering. Using the FindAllMarkers function of the Seurat package, we identified genes with log2 fold change >0.25 and adjusted P value >0.01 for each cluster. Based on the order of log2 fold change, the top 100 genes were further identified as markers of each cluster. To identify the genes specifically expressed in each cluster, gene markers that were present in at least three clusters were selected. We performed cluster analysis on GABAergic neurons by selecting the cells that express at least one of three markers of GABAergic neuron cells (Slc32a1, Gad1, and Gad2) in previously identified GABAergic neurons and clustered the remaining neuron cells using the FindAllClusters function in Seurat with the resolution value set as equal to 0.8. Differential expression genes between two sexes and biological status were obtained by using the FindMarkers function in Seurat.

## QUANTIFICATION AND STATISTICAL ANALYSIS

In experiments with paired samples, we used a paired t-test and a Wilcoxon matched-pairs signed-rank test. In all other experiments, we used a t-test, a t-test with Welch’s correction for unequal standard deviation, ANOVA for parametric data, and a Mann-Whitney U-test. All values are calculated with Prism 6.0 or MATLAB and presented as the mean ± SEM. All data are expressed as the mean ± SEM unless stated otherwise. In some of the plots, outlier values are shown as plus symbols. Data were processed and analyzed using MATLAB and GraphPad PRISM 8 (GraphPad Software). The sample sizes were chosen based on common practice in animal behavioral experiments. All data were analyzed with two-tailed tests. In the experiments with paired samples, we used the Wilcoxon matched-pairs signed-rank test. In the experiments with non-paired samples, we used the Mann–Whitney U-test or unpaired t-test with Welch’s correction for comparison between two groups and used the Kruskal-Wallis test with Dunn’s multiple comparisons correction, one-way ANOVA with Dunnett’s multiple comparisons test or with Tukey’s multiple comparisons test, two-way ANOVA for comparison among multiple groups. The significance threshold was held at α = 0.05, two-tailed (p > 0.05; *p < 0.05; **p < 0.01; ***p < 0.001. n.s, not significant). Full statistical analyses corresponding to each dataset are presented in Table S1 and Table S2.

---

**Extended Data Fig. 1.**
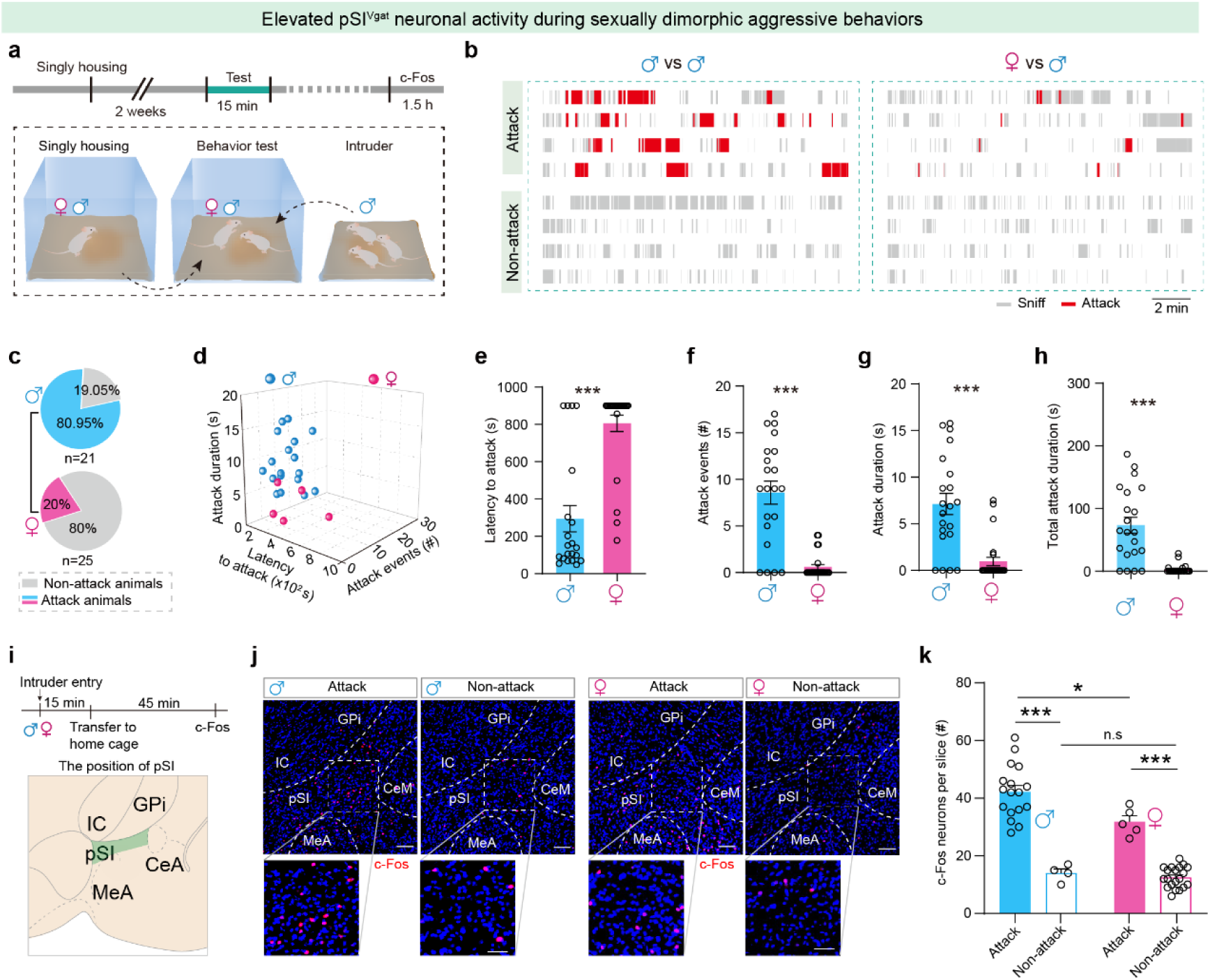
pSI neurons were selectively activated during male and female aggressive behavior. (**a**) Experimental scheme for the examination of aggressive behaviors and c-Fos expression in the pSI of male and female mice after resident-intruder test. **(b)** Raster plots of recorded social behaviors during 15-minute encounters when males and females are exposed to a male intruder (red blocks indicate the duration of episodes of attack, and grey blocks indicate the duration of episodes of sniff). (**c**) The probabilities of male and female mice that show attack. (**d**) Scatter plot of aggressive behaviors, including latency to the first attack, attack events, and attack duration, in all males (n = 21) and female (n = 25) mice. **(e-h)** Aggressive behaviors, including latency to the first attack **(e)**, attack events (f), attack duration (g) and total attack duration (h) in all males (n = 21) and female (n = 25) mice. **(i)** Illustrations adapted from the Allen Mouse Brain Atlas. GPi, globus pallidus internus; IC, internal capsule; CeM, central amygdaloid nucleus; MeA, medial amygdaloid nucleus. (**j**) Representative images of c-Fos^+^ neurons analysis in pSI after sniff (non-attack) or after attack in males and females. Scale bars represent 100 µm (upper) and 50 µm (lower). (**k**) Summary data of the number of c-Fos^+^ neurons in pSI after a behavioral paradigm of attack test. Data are mean ± SEM; (e) to (h), and (k): Mann-Whitney U test; **P < 0.01, ***P < 0.001, n.s, not significant. Statistical details are available in the supplementary table S1.

**Extended Data Fig. 2.**
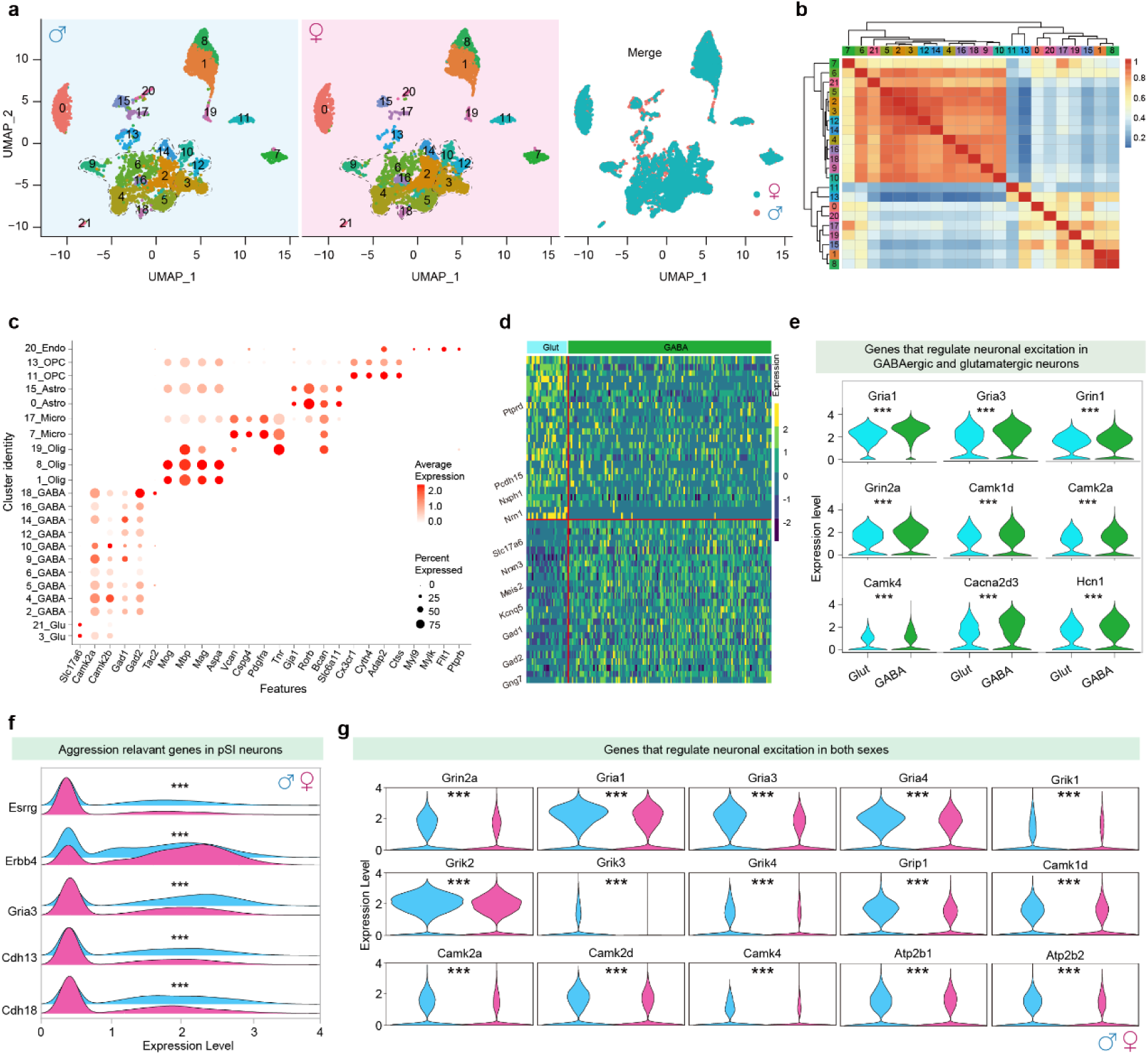
Single-nucleus RNA-seq of pSI neurons from male and female mice. (**a**) Two-dimensional UMPA visualization showing the distribution of cells from males (left), cells from females (middle), and merged cells (right) among major pSI cell types. (**b**) Analysis of the correlation of differentially expressed genes in neuronal clusters (**c**) Bubble chart of Z-scores of normalized expressions for selecting marker genes within all major cell types. (**d**) Differential genes of the GABAergic and glutamatergic cells from all pSI neurons. (**e**) Differential expression of the neuronal excitation genes in the GABAergic and glutamatergic cells. Violin plots show that the expression level of HCN1 genes with significantly higher in GABA neurons versus glutamatergic neurons in the pSI. (**f**) The volcano plot of aggression-relevant genes of males versus females in the pSI. **(g)** The violin plot of genes that regulate neuronal excitation of males versus females in the pSI. In (d) to (g): Wilcoxon rank-sum test; ***P < 0.001. For all panels, data are shown as the mean±SEM. Statistical details are available in the supplementary table S2.

**Extended Data Fig. 3.**
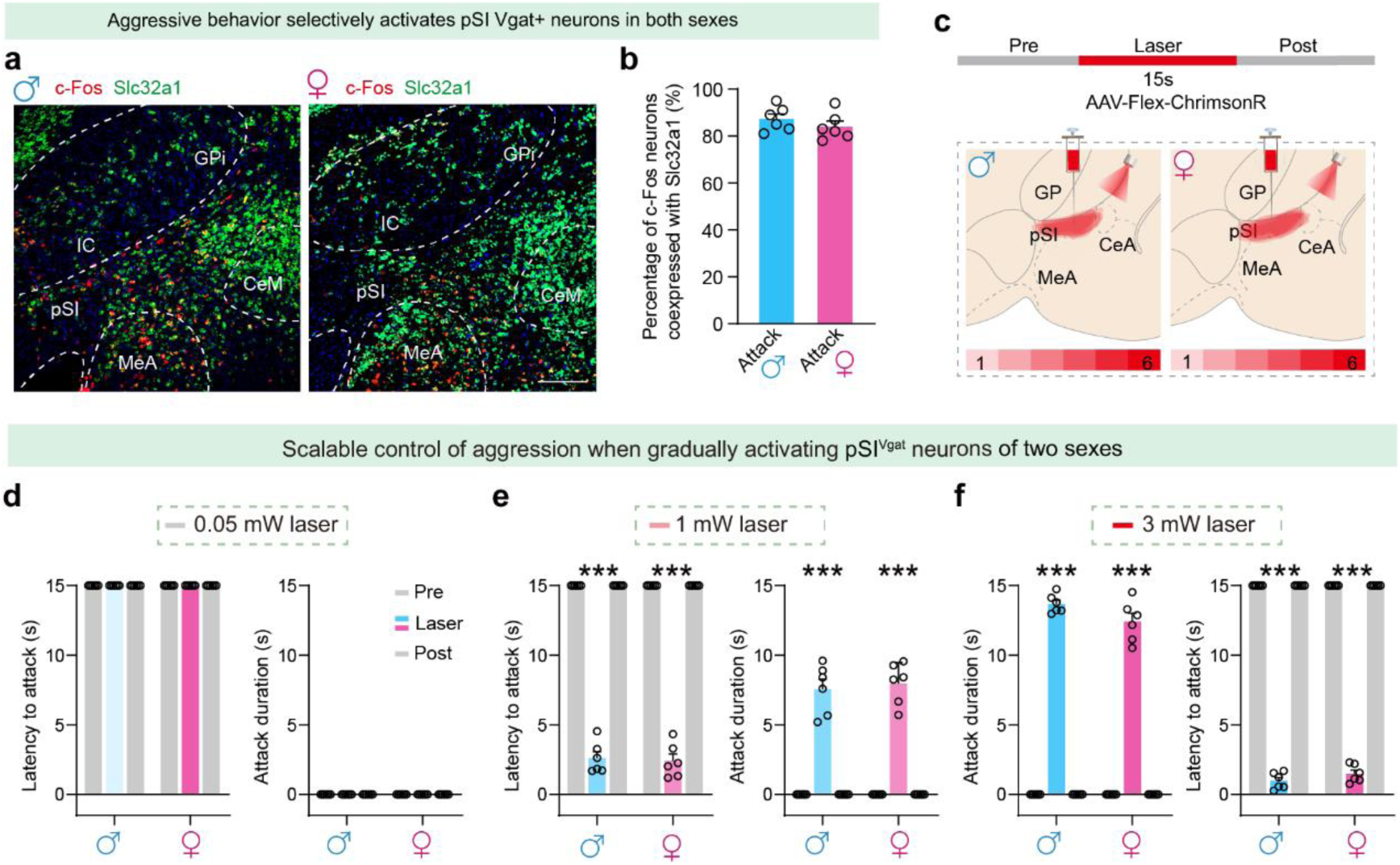
Neural dynamics of pSI^Vgat^ neurons is functionally correlated with aggression. (**a**) C-Fos and Slc32a1 co-expression in the pSI after aggressive behaviors in males (left) and females (right). Scale bar, 100 μm. (**b**) C-Fos ratio that co-expressed with Vgat in the pSI after aggressive behaviors in males and females. **(c)** Representative ChrimsonR signals are mapped onto coronal mouse brain atlas images in Vgat-Cre male and female mice. **(d-f)** Attack latency and attack duration with the graded poststimulation of 0.05 mW (d), 1 mW (e), and 3 mW (f) in Vgat-Cre males and females (n = 8 mice in each group). In (d) to (f): one-way ANOVA followed by post hoc Tukey’s multiple comparisons; ***P < 0.001, n.s, not significant. Statistical details are available in the supplementary table S1. Data are mean ± SEM.

**Extended Data Fig. 4.**
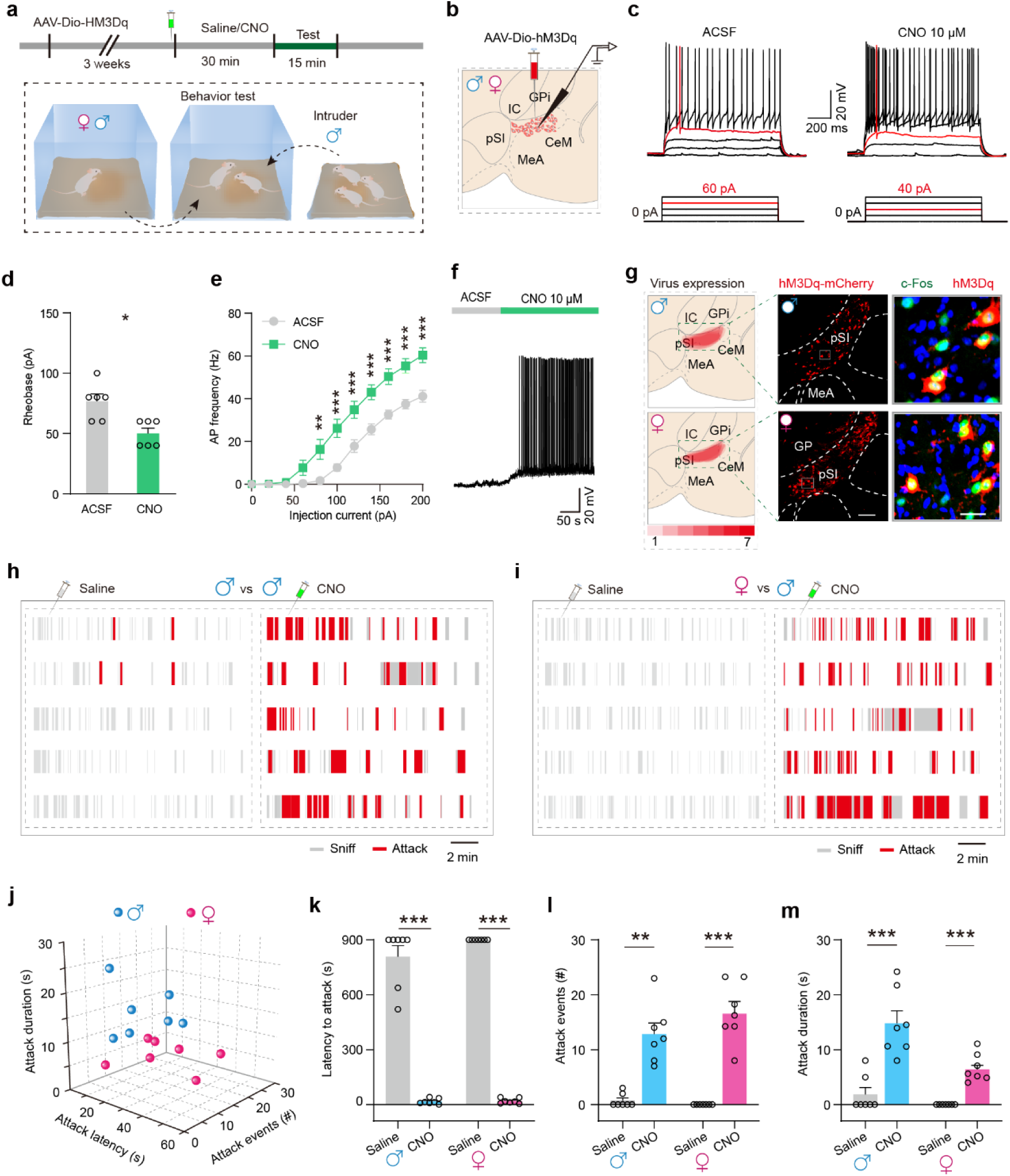
Pharmacogenetic activation of pSI^Vgat^ neurons triggers aggression. (**a**) Schematic of the viral injection strategy for the expression of AAV-Dio-hM3Dq-mCherry (red) in the pSI of Vgat-Cre male and female mice. **(b)** Schematic of whole-cell patch clamp recordings from hM3Dq-expressing pSI^Vgat^ neurons in Vgat-Cre male and female mice. (**c**) Representative spiking patterns were recorded in hM3Dq-expressing pSI^Vgat^ neurons in response to perfusion with CNO. Each injection train lasted for a duration of 1 s, with a step increase of 20 pA. (**d-e**) Comparison of rheobase (d, minimum activation current of the first AP), and AP frequency (e) in response to step current injections before or after treatment with CNO (n = 6 neurons from 2 mice). (**f**) Whole-cell patch clamp recording trace from hM3Dq-expressing pSI^Vgat^ neurons in response to perfusion with CNO. (**g**) Expression of AAV-Dio-hM3Dq-mCherry (red) in the pSI of Vgat-Cre male and female mice. (**h-i**) Example social behaviors after pharmacogenetic activation of pSI^Vgat^ neurons with saline or CNO in Vgat-Cre male and female mice. (**j-m**) Aggressive behaviors (j), including latency to the first attack (k), attack events (l), and attack duration (m) when pharmacogenetic activation of pSI^Vgat^ neurons in Vgat-Cre male and female mice. Data are mean ± SEM; (f): Wilcoxon matched-pairs signed rank test; (g): Two-way ANOVA followed by post-hoc Šidák’s multiple comparisons; (k) to (m): Paired t test; *P < 0.05, **P < 0.01, ***P < 0.001. Statistical details are available in the supplementary table S1.

**Extended Data Fig. 5.**
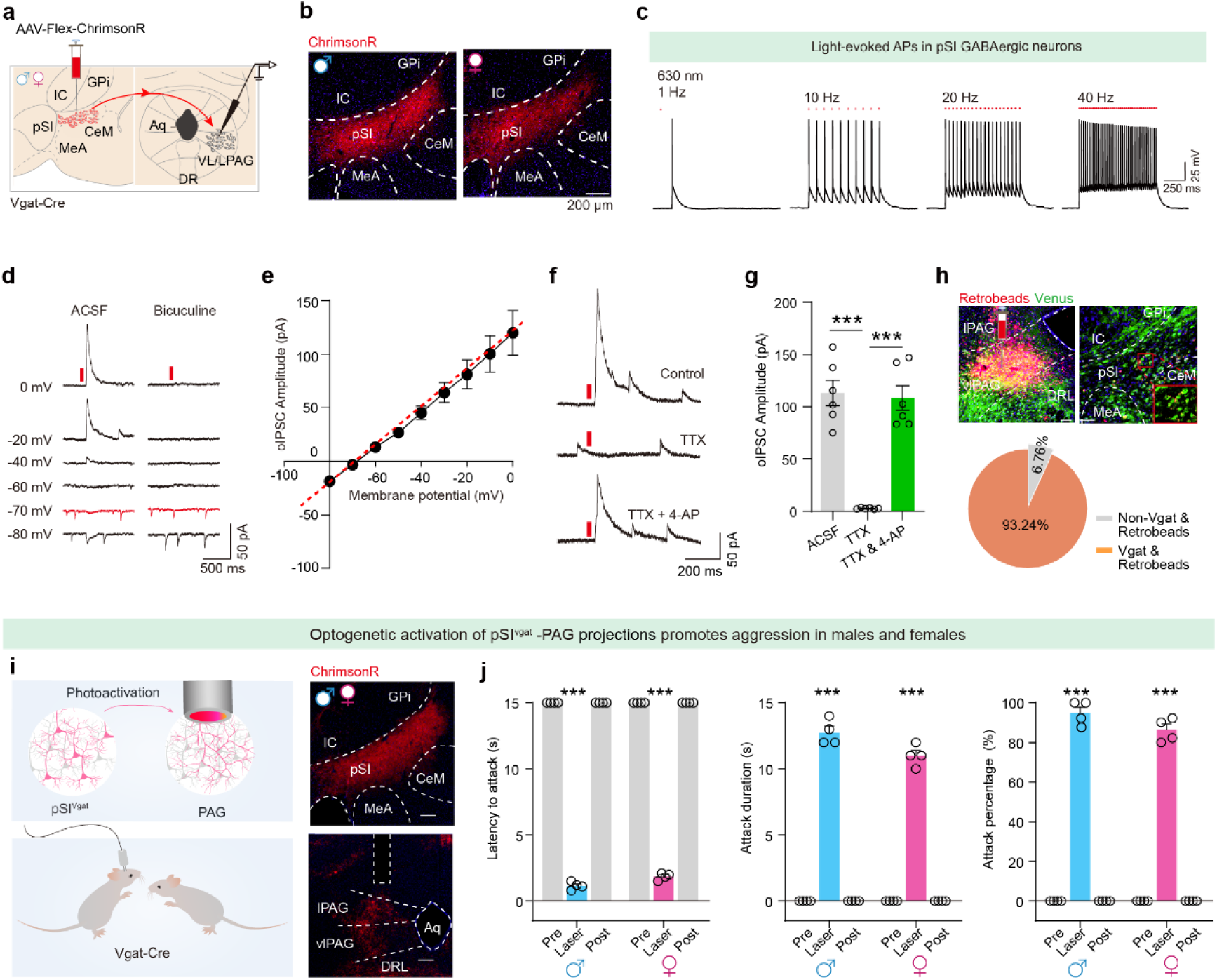
Optogenetic activation of pSI^Vgat^-vl/lPAG projection terminals drives time-locked aggressive behavior in both sexes. (**a**) Schematic of whole-cell patch clamp recordings from pSI^Vgat^ neurons and vl/lPAG neurons in male and female mice. (**b**) Representative images of pSI neurons infected with AAV_2/9_-Dio-ChrimsonR (scale bars, 200 μm) in Vgat-Cre mice. (**c**) Sample recording trace from a ChrimsonR-expressing pSI neuron in response to light-pulse trains (pulse: 638 nm, 5 ms, 40 Hz) evoked phase-locked spiking activity. (**d**) Representative traces of the light-evoked postsynaptic currents in the vl/lPAG neurons by photostimulating (red laser, 5 mW) axonal projections from the pSI^Vgat^ to vl/lPAG neurons in the absence or presence of the GABA receptor antagonist bicuculine (100 μM); the membrane potential holding at − 80 mV, − 70 mV, − 60 mV, − 40 mV, − 20 mV, and 0 mV, respectively. (**e**) Line plot of the amplitudes of light-evoked postsynaptic currents based on diverse membrane potentials (n=6 cells). According to the linear relationship, calculated the reversal potential is about − 67.69 mV. (**f-g**) Representative traces (f) and amplitude (g) of oIPSCs evoked by photostimulation in male mice with or without bath-application of TTX (1 μM) and 4-AP (100 μM). **(h)** Upper, representative images show infections of retrobeads in the PAG, and the retrograde labeling of pSI^-PAG^ neurons (red) was overlaid with GABAergic pSI neuros (green) in Vgat-venus mice. Lower, the ratio of pSI^-PAG^ neurons (red) that co-expressed with Vgat-venus (green) in the pSI. (**i-j**) Behavioral paradigm (i) and summary data (j) showing a socially housed male or female mouse in a novel cage initiating attacks on a socially housed male intruder after optical activation of the pSI^Vgat^-PAG projection. Attack latency (left), attack duration (middle), and attack percentage (right) induced by optogenetics during the pre-laser, laser, and post-laser phases in the male (n = 4 mice) and female (n = 4 mice) mice. Data are mean ± SEM; (g): One-way ANOVA followed by Dunnett’s multiple comparisons test; (j): One-way ANOVA followed by Tukey’s multiple comparisons test; ***P < 0.001. Statistical details are available in the supplementary table S1.

**Extended Data Fig. 6.**
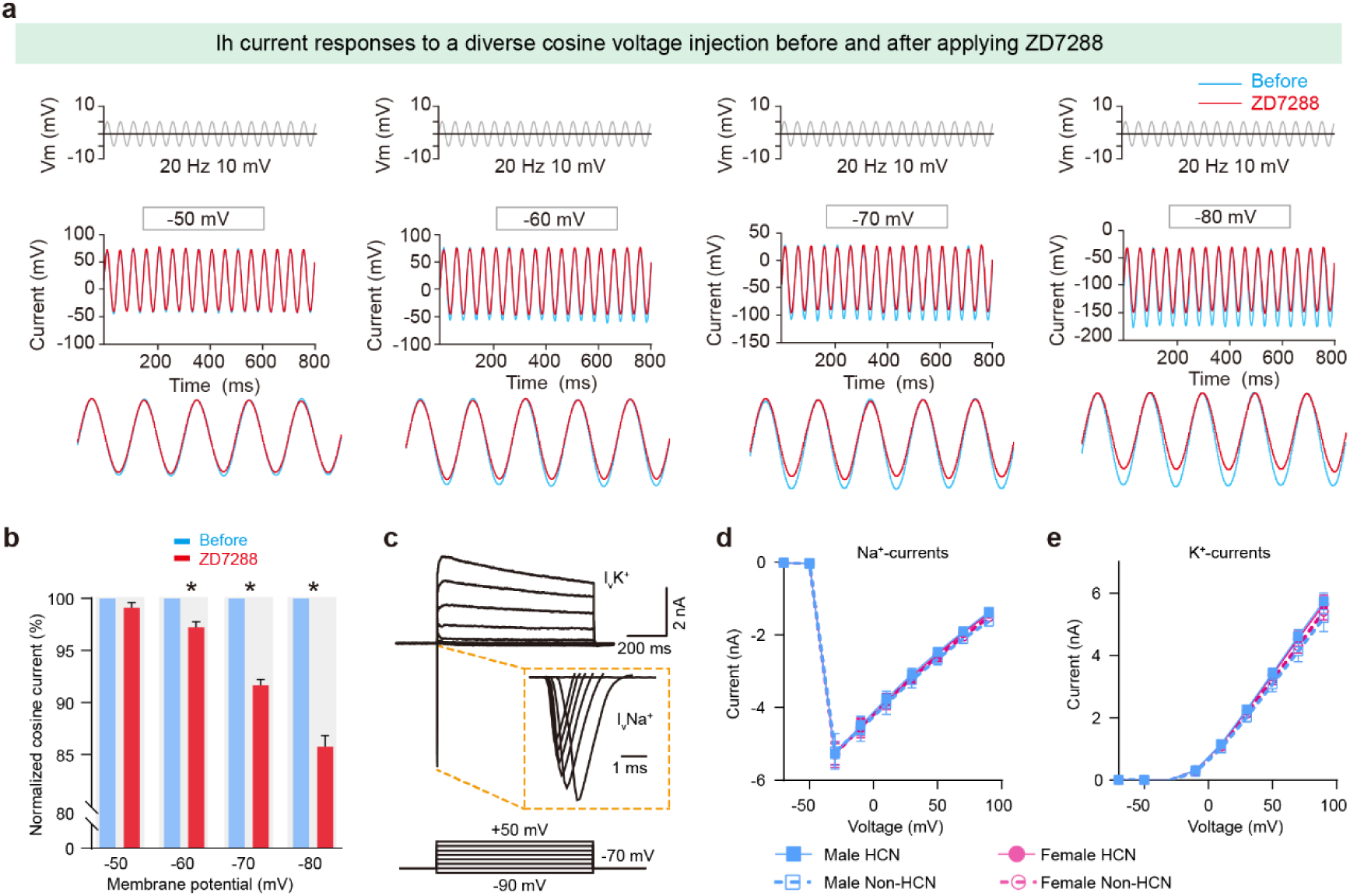
Ih current regulates the excitability of pSI^-PAG^ neurons in male and female mice. (**a**) Upper, experimental protocol of a diverse chirp stimulus. Lower, example current responses to a diverse chirp stimulus voltage injection (− 50 mV, − 60 mV, − 70 mV, − 80 mV) in pSI^-PAG^ HCN neurons. (**b**) Summarized data of normalized cosine currents in pSI^-PAG^ neurons recorded from HCN neurons before or after treatment with ZD7288 (100 μM). (**c**) Experimental protocol and sample traces recorded from pSI^-PAG^ neurons in the voltage clamp mode. (**d-e**) Summarized data of sodium current (Na^+^, g) and potassium current (K^+^, h) in pSI^-PAG^ neurons recorded from HCN neurons or non-HCN neurons in male and female mice. Data are mean ± SEM; in (b), Wilcoxon matched-pairs signed rank test; (d) and (e): Two-way ANOVA followed by *Post-hoc* Šidák’s multiple comparisons; *P < 0.05, **P < 0.01, ***P < 0.001, n.s, not significant. Statistical details are available in the supplementary table1.

**Extended Data Fig. 7.**
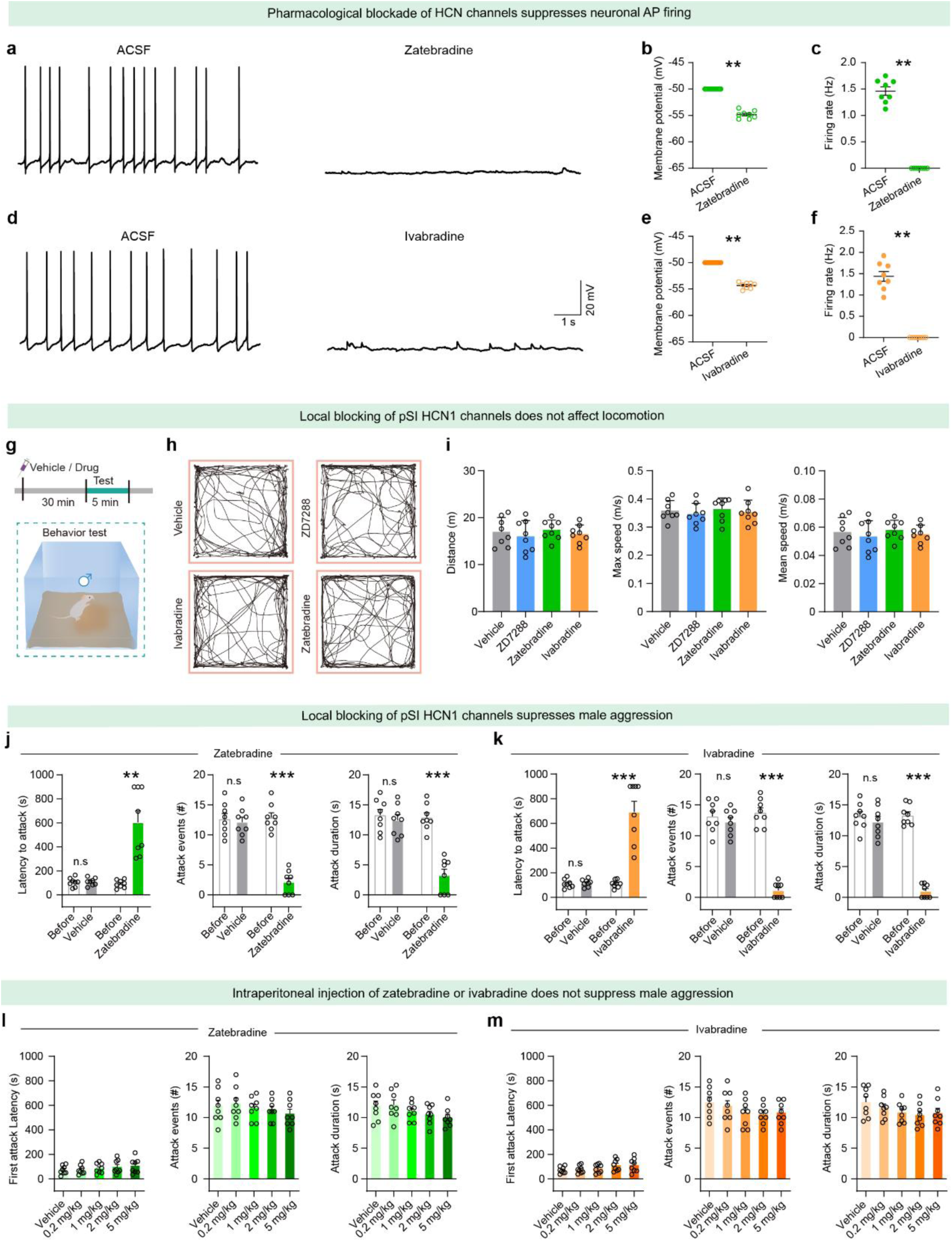
Pharmacological manipulation of HCN1 channels alters male aggressive behaviors. **(a-c)** Example electrophysiological traces, RMP, and firing rate of pSI neurons with pharmacological inhibition of HCN1 channels in the pSI by zatebradine. **(d-f)** Example electrophysiological traces, RMP, and firing rate of pSI neurons with pharmacological inhibition of HCN1 channels in the pSI by ivabradine. **(g-i)** Upper, timeline of the animal model of male for a vehicle or ZD7288, zatebradine, and ivabradine. Lower, the behavioral paradigm during the open field test treatment with HCN channel bolckers (n = 8 mice in each group). Statistical data showing the distance, mean speed, and max speed in the 5-minute test. Significance was determined by the Mann-Whitney U test. **(j-k)** Effects of pharmacological inhibition of HCN1 channels (i.p. injection of the zatebradine, and ivabradine) during the attack test for single-housed male mice toward a male intruder in its home cage. Attack latency, attack events, and attack duration of male mice, in an administered intraperitoneally manner. **(l-m)** Aggressive behaviors for inhibiting HCN1 channels with pharmacological inhibition of HCN1 channels in the pSI by vehicle control, ZD7288, zatebradine, and ivabradine during the aggression test (left, n = 8 mice in each group). Statistical data showing attack latency, attack duration, and attack events. Data are mean ± SEM; (b), (c), (e), and (f): Wilcoxon matched-pairs signed rank test; (i), Unpaired t test; (j) and (k), Paired t test; (l) and (m), One way ANOVA followed by Holm-Šídák’s multiple comparisons test; **P < 0.01, ***P < 0.001, n.s, not significant. Statistical details are available in the supplementary table S1.

**Extended Data Fig. 8.**
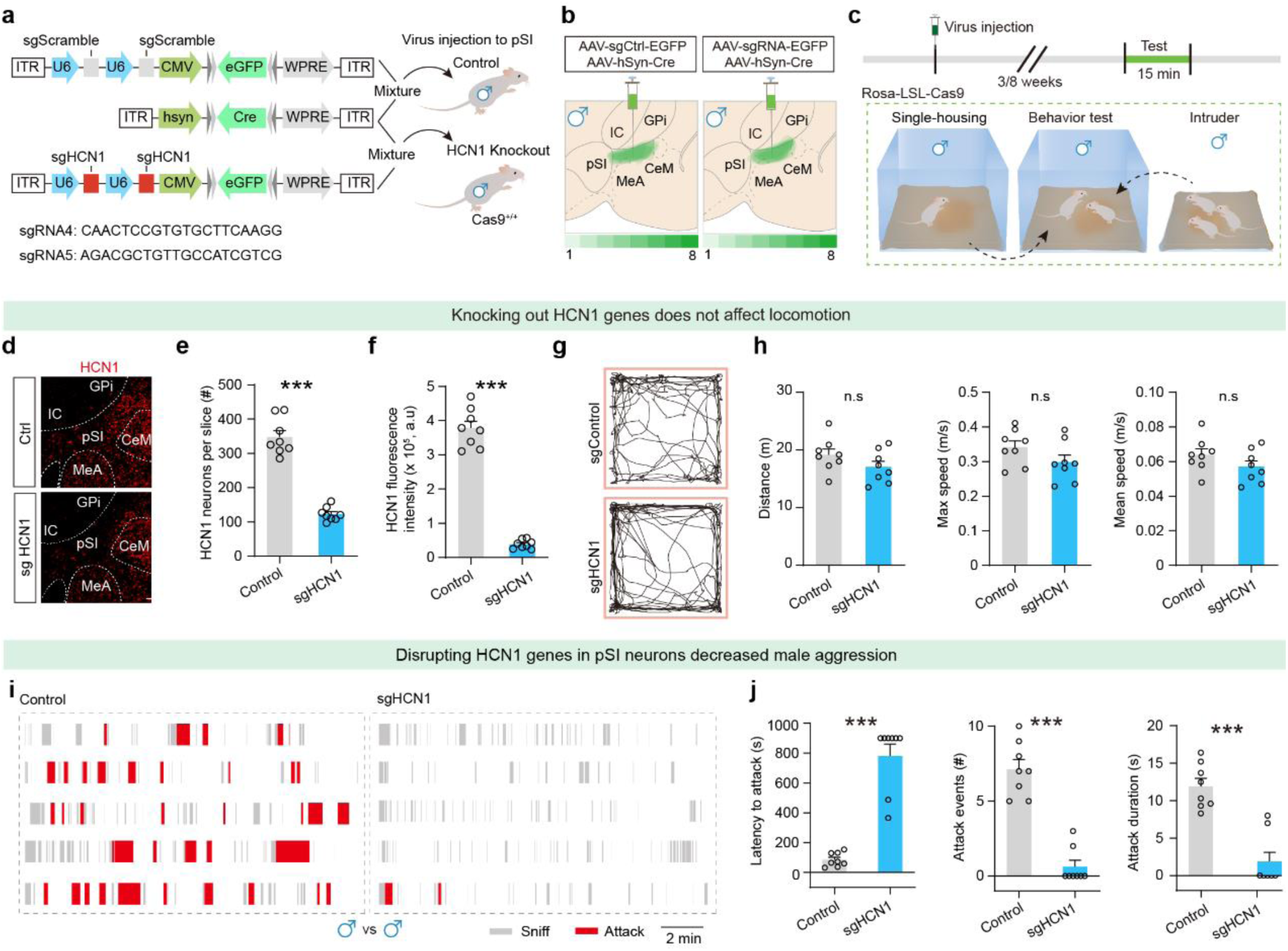
Knockout of HCN1 genes in pSI neurons suppress attacks in males. (**a-c**) Design of examining the knockout of HCN1 genes in GABAergic pSI neurons suppress attacks in males with the sgHCN1 genes (a), injection (b), and behavioral paradigm (c). **(d**) Representative slices showing HCN1 expression in neurons of the pSI. **(e-f)** Total HCN1 positive cells (b), and fluorescence intensity of positive cells (c, green) in the pSI of females. **(g)** Example trajectory trace for sgControl and sgHCN1 group to examine the open field test (left, n = 8 mice per group). **(h)** Statistical data showing the distance, mean speed, and max speed in the 5-minute test. (**i**) Raster plots of attack behavior recorded during 15-minute encounters in the sgControl and sgHCN1 groups (red blocks indicate the duration of episodes of attack, and gray blocks indicate the duration of episodes of sniff, n = 8 mice per group). **(j)** Latency to the first attack, attack events, and attack duration at 8 weeks after the viral injection, as illustrated in (a). Data are mean ± SEM; in (e), (f), (h), and (j): Unpaired t test; ***P < 0.001, n.s, not significant. Statistical details are available in the supplementary table S1.

**Extended Data Fig. 9.**
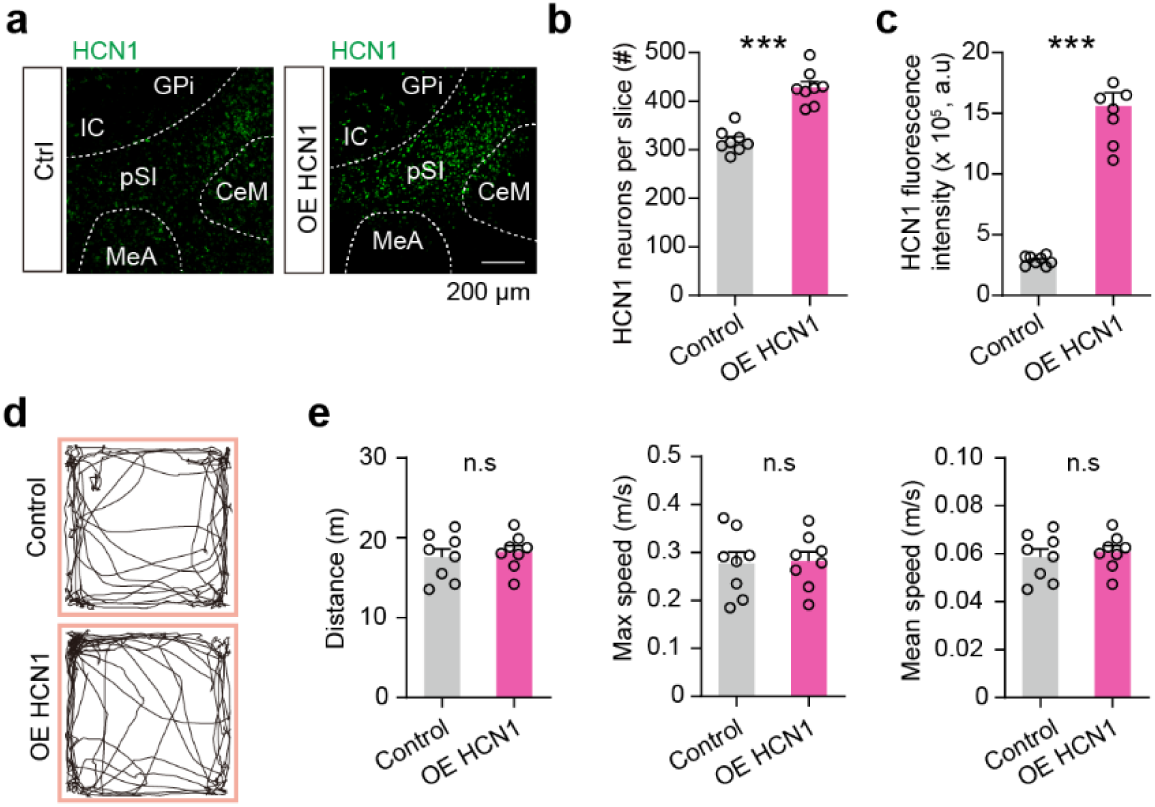
Overexpressing of HCN1 genes in GABAergic pSI neurons did not affect locomotion. **(a)** Representative slices showing HCN1 expression in neurons of the pSI. **(b-c)** Total HCN1 positive cells (b), and fluorescence intensity of positive cells (c, green) in the pSI of females. **(d)** Example trajectory trace for the control and overexpression of HCN1 groups to examine the open field test (left, n = 8 mice/group). **(e)** Statistical data showing the distance, mean speed, and max speed in the 5-minute test. Data are mean ± SEM; (b), (c), and (e): Unpaired t test; ***P < 0.001, n.s, not significant. Statistical details are available in the supplementary table S1.

**Extended Data Fig. 10.**
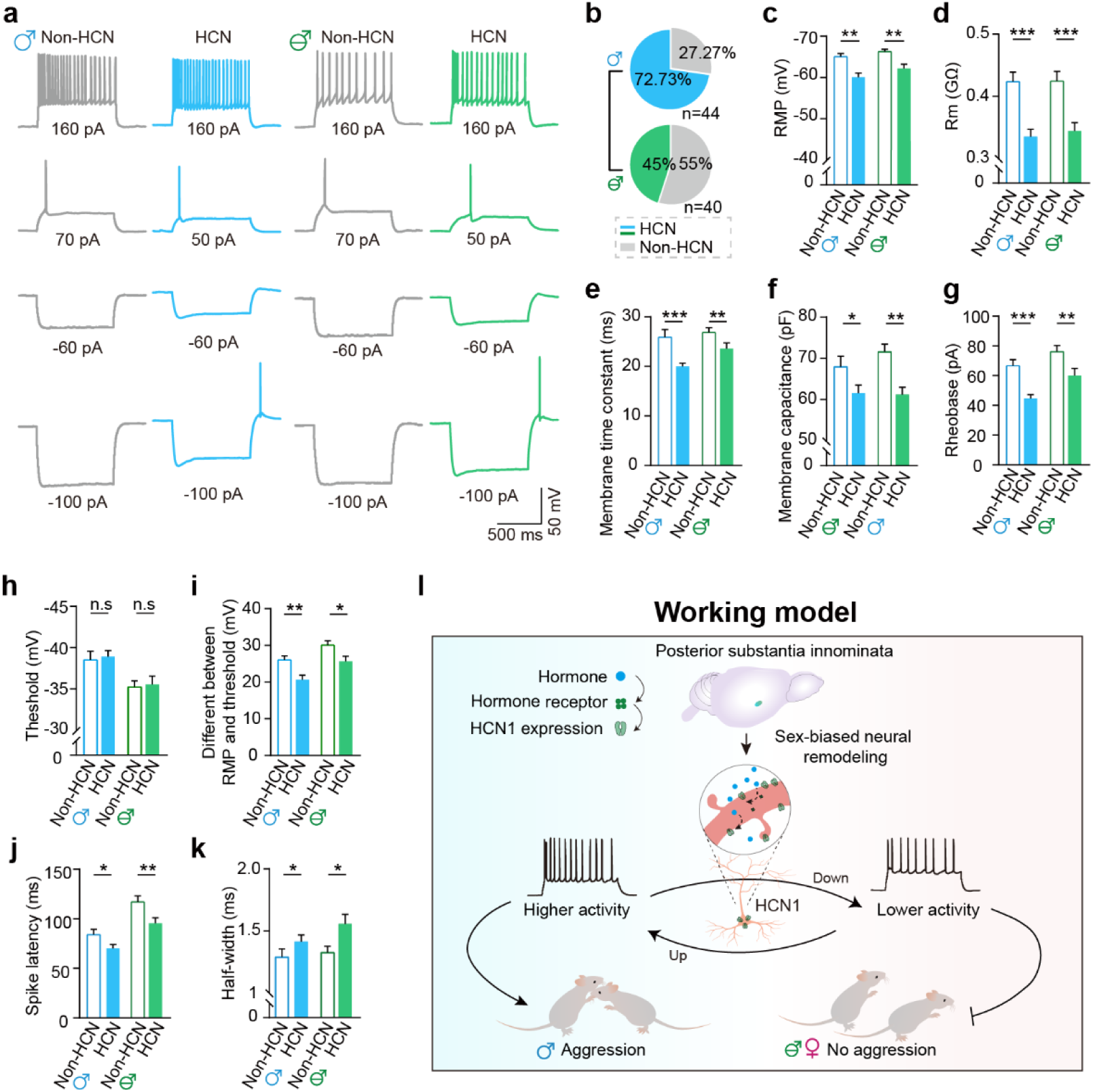
Electrophysiological properties of pSI^-PAG^ neurons in male and castrated male mice. (**a**) Representative spiking patterns of pSI^-PAG^ neurons in male and castrated male mice. Example voltage signals recorded from a pSI^-PAG^ neuron from the male and castrated male mice in response to intracellular current injection of a pulse train from -100 pA to 160 pA in 10 pA steps for 1 s. (**b**) Relative proportions of HCN neurons (green) and non-HCN (gray) pSI^-PAG^ neurons in male (upper, n = 44 neurons from 5 males) and castrated male (lower, n = 40 neurons from 5 castrated males) mice. (**c-k**) Comparison of basic electrophysiological properties of pSI^-PAG^ neurons between male and castrated male mice. (c) RMP, (d) membrane resistance, (e) membrane time constant, (f) membrane capacitance of all the recorded pSI-PAG neurons. Comparison of other parameters including (g) rheobase, (h) AP threshold, (i) difference between RMP and AP threshold, (j) spike latency, and (k) half-width between pSI^-PAG^ neurons (male, n = 44 neurons versus castrated male, n = 40 neurons from 10 mice each group). **(l)** Schematics for the HCN1-mediated hyperexcitability for mediating heightened aggression. Data are mean ± SEM; (b) to (k): Mann-Whitney U test; *P< 0.05, **P < 0.01, ***P < 0.001, n.s, not significant. Statistical details are available in the supplementary table S1.

## Key resources table

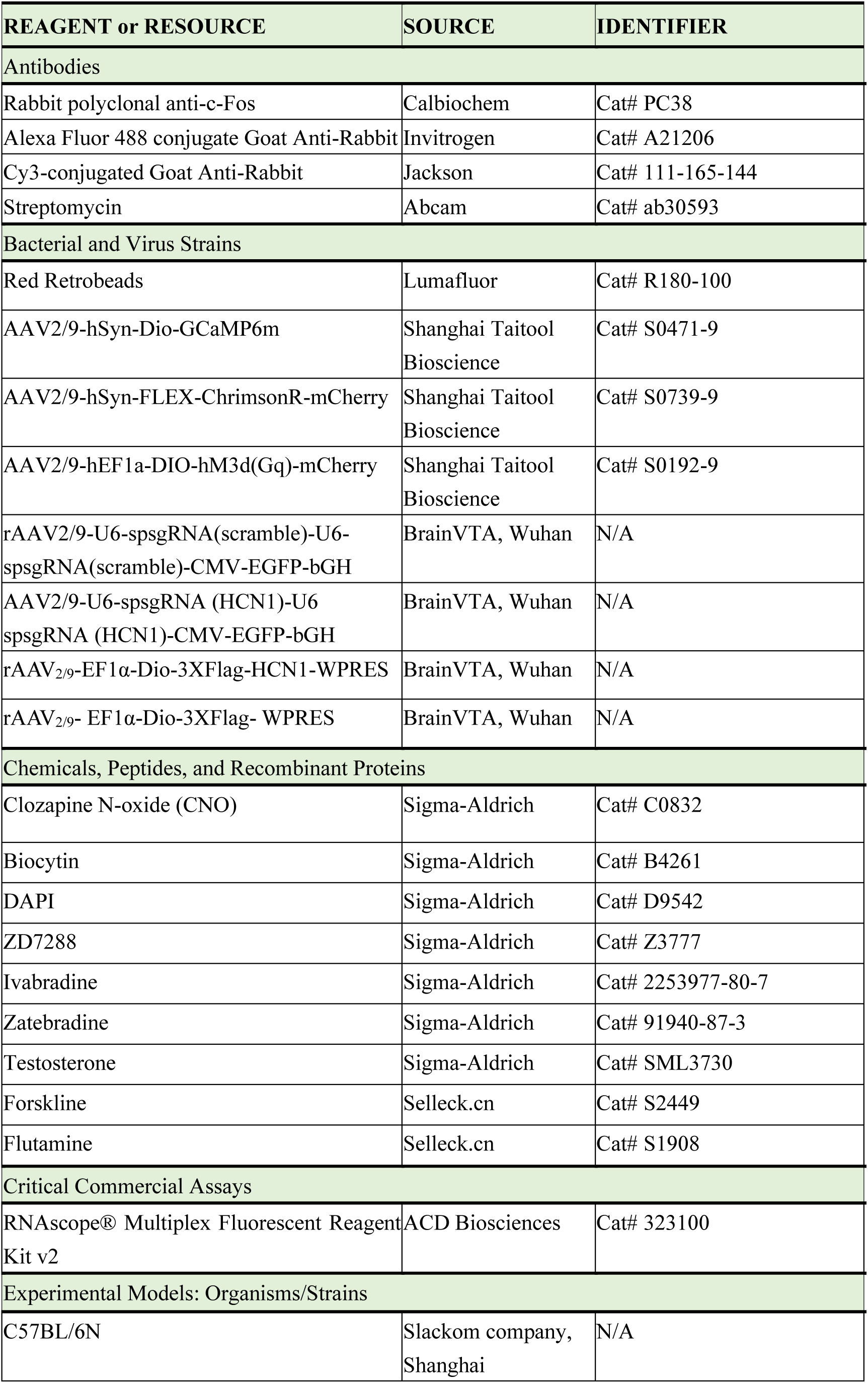
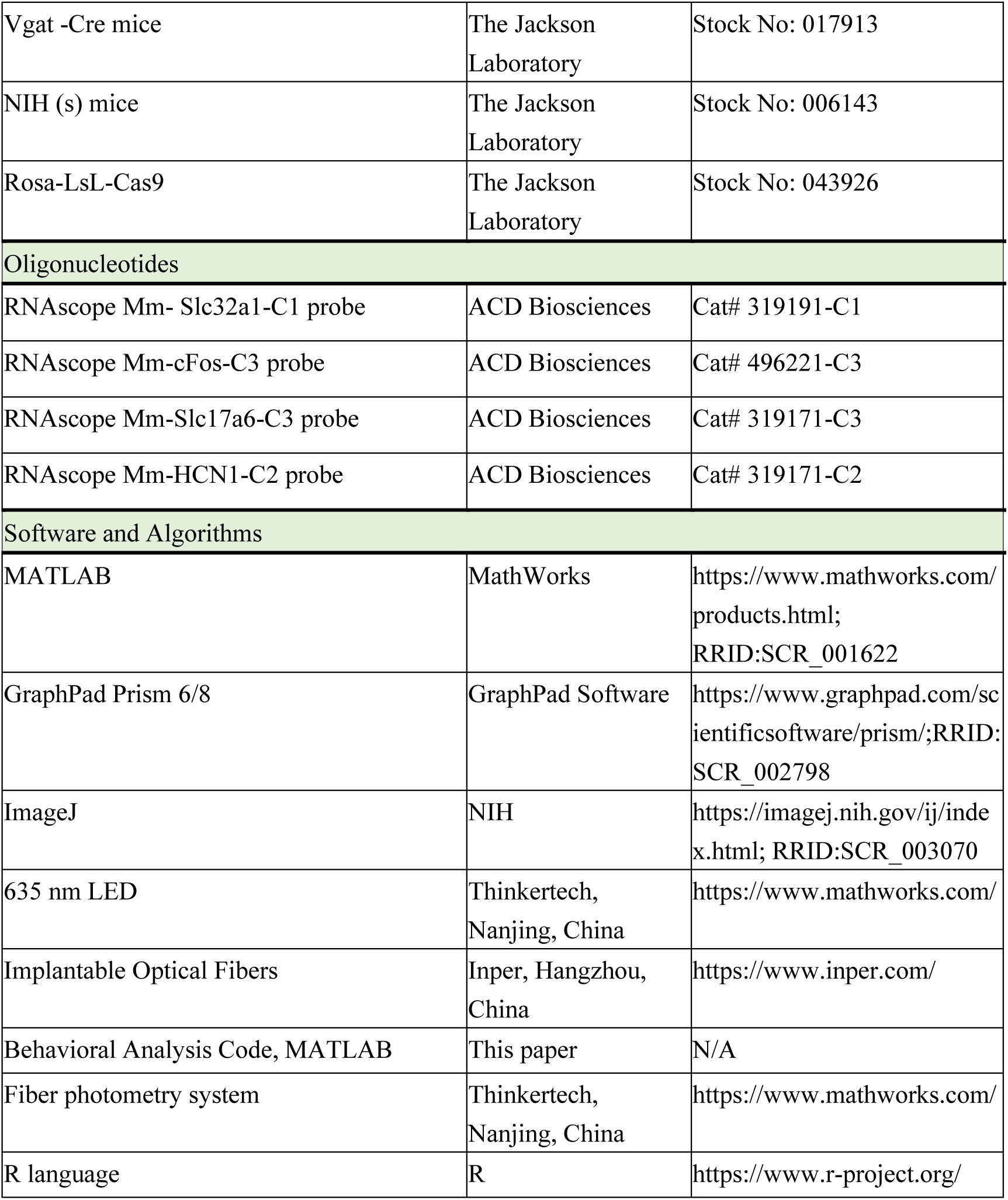

## References

1. C. F. Yang, N. M. Shah, Representing sex in the brain, one module at a time. Neuron 82, 261–278 (2014).

2. K. Hashikawa, Y. Hashikawa, J. Lischinsky, D. Lin, The Neural Mechanisms of Sexually Dimorphic Aggressive Behaviors. Trends Genet 34, 755–776 (2018).

3. J. A. Tobias, R. Montgomerie, B. E. Lyon, The evolution of female ornaments and weaponry: social selection, sexual selection and ecological competition. Philos Trans R Soc Lond B Biol Sci 367, 2274–2293 (2012).

4. Z. Zhu et al., A substantia innominata-midbrain circuit controls a general aggressive response. Neuron 109, 1540–1553 e1549 (2021).

5. E. K. Unger et al., Medial amygdalar aromatase neurons regulate aggression in both sexes. Cell Rep 10, 453–462 (2015).

6. K. Hashikawa et al., Esr1(+) cells in the ventromedial hypothalamus control female aggression. Nat Neurosci 20, 1580–1590 (2017).

7. Z. Zhu et al., A hypothalamic-amygdala circuit underlying sexually dimorphic aggression. Neuron, (2024).

8. P. Chen, W. Hong, Neural Circuit Mechanisms of Social Behavior. Neuron 98, 16–30 (2018).

9. T. Yang, N. M. Shah, Molecular and neural control of sexually dimorphic social behaviors. Curr Opin Neurobiol 38, 89–95 (2016).

10. C. Dulac, T. Kimchi, Neural mechanisms underlying sex-specific behaviors in vertebrates. Curr Opin Neurobiol 17, 675–683 (2007).

11. B. N. Dugger, J. A. Morris, C. L. Jordan, S. M. Breedlove, Androgen receptors are required for full masculinization of the ventromedial hypothalamus (VMH) in rats. Horm Behav 51, 195–201 (2007).

12. M. V. Wu et al., Estrogen masculinizes neural pathways and sex-specific behaviors. Cell 139, 61–72 (2009).

13. S. A. Juntti et al., The androgen receptor governs the execution, but not programming, of male sexual and territorial behaviors. Neuron 66, 260–272 (2010).

14. C. F. Yang et al., Sexually dimorphic neurons in the ventromedial hypothalamus govern mating in both sexes and aggression in males. Cell 153, 896–909 (2013).

15. H. Lee et al., Scalable control of mounting and attack by Esr1+ neurons in the ventromedial hypothalamus. Nature 509, 627–632 (2014).

16. R. Ammari et al., Hormone-mediated neural remodeling orchestrates parenting onset during pregnancy. Science 382, 76–81 (2023).

17. B. Gegenhuber, M. V. Wu, R. Bronstein, J. Tollkuhn, Gene regulation by gonadal hormone receptors underlies brain sex differences. Nature 606, 153–159 (2022).

18. F. Li et al., Sex differences orchestrated by androgens at single-cell resolution. Nature 629, 193–200 (2024).

19. S. B. Li et al., Hyperexcitable arousal circuits drive sleep instability during aging. Science 375, eabh3021 (2022).

20. M. F. Nolan et al., A behavioral role for dendritic integration: HCN1 channels constrain spatial memory and plasticity at inputs to distal dendrites of CA1 pyramidal neurons. Cell 119, 719–732 (2004).

21. M. F. Nolan et al., The hyperpolarization-activated HCN1 channel is important for motor learning and neuronal integration by cerebellar Purkinje cells. Cell 115, 551–564 (2003).

22. S. Vesuna et al., Deep posteromedial cortical rhythm in dissociation. Nature 586, 87–94 (2020).

23. X. Zhou et al., Hyperexcited limbic neurons represent sexual satiety and reduce mating motivation. Science 379, 820–825 (2023).

24. T. Kimchi, J. Xu, C. Dulac, A functional circuit underlying male sexual behaviour in the female mouse brain. Nature 448, 1009–1014 (2007).

25. V. S. Mandiyan, J. K. Coats, N. M. Shah, Deficits in sexual and aggressive behaviors in Cnga2 mutant mice. Nat Neurosci 8, 1660–1662 (2005).

26. L. Stowers, T. E. Holy, M. Meister, C. Dulac, G. Koentges, Loss of sex discrimination and male-male aggression in mice deficient for TRP2. Science 295, 1493–1500 (2002).

27. A. Ludwig, X. Zong, M. Jeglitsch, F. Hofmann, M. Biel, A family of hyperpolarization-activated mammalian cation channels. Nature 393, 587–591 (1998).

28. B. Santoro et al., Identification of a gene encoding a hyperpolarization-activated pacemaker channel of brain. Cell 93, 717–729 (1998).

29. D. DiFrancesco, P. Tortora, Direct activation of cardiac pacemaker channels by intracellular cyclic AMP. Nature 351, 145–147 (1991).

30. A. Porro et al., A high affinity switch for cAMP in the HCN pacemaker channels. Nat Commun 15, 843 (2024).

31. A. Porro et al., The HCN domain couples voltage gating and cAMP response in hyperpolarization-activated cyclic nucleotide-gated channels. Elife 8, (2019).

32. M. V. Wu, N. M. Shah, Control of masculinization of the brain and behavior. Curr Opin Neurobiol 21, 116–123 (2011).

33. D. J. Anderson, Circuit modules linking internal states and social behaviour in flies and mice. Nat Rev Neurosci 17, 692–704 (2016).

34. H. Chiu et al., A circuit logic for sexually shared and dimorphic aggressive behaviors in Drosophila. Cell 184, 847 (2021).

35. M. Zelikowsky et al., The Neuropeptide Tac2 Controls a Distributed Brain State Induced by Chronic Social Isolation Stress. Cell 173, 1265–1279 e1219 (2018).

36. T. Yang et al., Social Control of Hypothalamus-Mediated Male Aggression. Neuron 95, 955–970 e954 (2017).

37. S. X. Zhang et al., Hypothalamic dopamine neurons motivate mating through persistent cAMP signalling. Nature 597, 245–249 (2021).

38. M. Liu, D. W. Kim, H. Zeng, D. J. Anderson, Make war not love: The neural substrate underlying a state-dependent switch in female social behavior. Neuron 110, 841–856 e846 (2022).

39. J. R. Moffitt et al., Molecular, spatial, and functional single-cell profiling of the hypothalamic preoptic region. Science 362, (2018).

40. P. B. Chen et al., Sexually Dimorphic Control of Parenting Behavior by the Medial Amygdala. Cell 176, 1206–1221 e1218 (2019).

41. G. Chen et al., Cellular and circuit architecture of the lateral septum for reward processing. Neuron, (2024).

42. C. S. Kim, P. Y. Chang, D. Johnston, Enhancement of dorsal hippocampal activity by knockdown of HCN1 channels leads to anxiolytic- and antidepressant-like behaviors. Neuron 75, 503–516 (2012).

43. C. Tsantoulas et al., Hyperpolarization-activated cyclic nucleotide-gated 2 (HCN2) ion channels drive pain in mouse models of diabetic neuropathy. Sci Transl Med 9, eaam6072 (2017).

44. P. Zhong et al., HCN2 channels in the ventral tegmental area regulate behavioral responses to chronic stress. Elife 7, (2018).

45. J. Cheng, G. Umschweif, J. Leung, Y. Sagi, P. Greengard, HCN2 Channels in Cholinergic Interneurons of Nucleus Accumbens Shell Regulate Depressive Behaviors. Neuron 101, 662–672 e665 (2019).

46. J. Lezmy et al., Astrocyte Ca(2+)-evoked ATP release regulates myelinated axon excitability and conduction speed. Science 374, eabh2858 (2021).

47. K. A. Lyman et al., Hippocampal cAMP regulates HCN channel function on two time scales with differential effects on animal behavior. Sci Transl Med 13, eabl4580 (2021).

48. J. W. McKinley et al., Dopamine Deficiency Reduces Striatal Cholinergic Interneuron Function in Models of Parkinson’s Disease. Neuron 103, 1056–1072 e1056 (2019).

49. M. Zhu et al., Shank3-deficient thalamocortical neurons show HCN channelopathy and alterations in intrinsic electrical properties. J Physiol 596, 1259–1276 (2018).

50. O. Postea, M. Biel, Exploring HCN channels as novel drug targets. Nat Rev Drug Discov 10, 903–914 (2011).

51. S. Stagkourakis, G. Spigolon, G. Liu, D. J. Anderson, Experience-dependent plasticity in an innate social behavior is mediated by hypothalamic LTP. Proc Natl Acad Sci U S A 117, 25789–25799 (2020).

52. S. Inoue et al., Periodic Remodeling in a Neural Circuit Governs Timing of Female Sexual Behavior. Cell 179, 1393–1408 e1316 (2019).

53. W. C. Krause et al., Oestrogen engages brain MC4R signalling to drive physical activity in female mice. Nature 599, 131–135 (2021).

